# Avoidance, confusion or solitude? Modelling how noise pollution affects whale migration

**DOI:** 10.1101/2023.09.27.559669

**Authors:** Stuart T. Johnston, Kevin J. Painter

## Abstract

Many baleen whales are renowned for their acoustic communication. Under pristine conditions, this communication can plausibly occur across hundreds of kilometres. Frequent vocalisations may allow a dispersed migrating group to maintain contact, and therefore benefit from improved navigation via the “wisdom of the crowd.” Human activities have considerably inflated ocean noise levels. Here we develop a data-driven mathematical model to investigate how ambient noise levels may inhibit whale migration. Mathematical models allow us to simultaneously simulate collective whale migration behaviour, auditory cue detection, and noise propagation. Rising ambient noise levels are hypothesised to influence navigation through three mechanisms: (i) diminished communication space; (ii) reduced ability to hear external sound cues and; (iii) triggering noise avoidance behaviour. Comparing pristine and current soundscapes, we observe navigation impairment that ranges from mild (increased journey time) to extreme (failed navigation). Notably, the three mechanisms induce qualitatively different impacts on migration behaviour. We demonstrate the model’s potential predictive power, exploring the extent to which migration may be altered under future shipping and construction scenarios.

## Introduction

Many whales routinely perform immense migrations [1], with individual whales observed travelling close to 20,000 km in a single year [2]. This clearly represents a significant investment of time and energy. The inherent difficulties of observing whale behaviour leaves numerous questions about navigation unanswered, not least the nature of navigation cues [1, 3]. One factor that has received considerable attention, though, is the whales’ ability to detect, respond to, and produce sounds [4]. Notably, sound propagates rapidly in water with little transmission loss, allowing information to be signalled/received across large distances [5]. External sound sources may provide navigating cues [3, 6], while echolocation may be used to probe the local environment [7].

Studies into “whalesong” that date back over half a century [8] have increased awareness of acoustic whale communication. In the context of navigation, this has been suggested to allow whales to broadcast and reinforce route information [5]. Such collective navigation has been investigated both experimentally and through modelling [9]. In the latter, collective behaviour has been shown to improve migration efficiency through a “many wrongs” principle [10] that reduces individual-level uncertainty, particularly if intrinsic navigation information is low [11, 12]. The many wrongs principle suggests that collective navigation is more effective due to the averaging out of navigational errors across the group [11]. Baleen whales can generate loud and low frequency sounds that lead to extraordinary communication ranges, theoretically covering hundreds of kilometres [4, 5]. As such, seemingly-isolated whales may still be benefiting from collective navigation via long-distance communication across a widely dispersed group [5, 13]. Substance to such conjectures can be found in the sequences of calls made by humpbacks, tens of kilometres apart, while traversing migration routes [14] or the apparent exchange of calls between bowhead whales as they navigate around ice [15].

The distance that a sound remains detectable in the ocean depends on fixed elements, such as bathymetry, and changing elements, such as ambient noise [16]. In the pre-industrial ocean, ambient noise would have been generated from factors such as wind, rain, breaking ice and biotic sources. Today, ambient noise in the ocean is increasingly a result of human activities [16, 17]. Anthropogenic sound sources include those from shipping, sonar, exploration, and offshore construction; estimates suggest that these may have already induced a rise of more than 20 dB in certain regions [16]. Given the importance of sound for communication and information, this noise pollution is believed to have a broad spectrum of impacts on whales (and other marine animals) [17–19]. This ranges from a reduction in communication range [5, 20] to physiological damage and stranding events that follow extreme noise events [19]. Observed reactions include altered swimming behaviour due to noise avoidance responses [21–23], and increased call volumes (Lombard effect) under higher ambient noise levels [24, 25]. Independent of any direct behavioural change, the communication range is still significantly diminished [24–26].

Mathematical models provide a framework to investigate the interplay between cue detection, noise pollution, and navigation behaviour. Abstract models have previously been developed to explore the benefit of collective navigation [9, 11, 12, 27, 28]. Broadly, we observe an increase in navigational efficiency with an increase in the number of observable conspecifics within a group; with specific investigations that include, for example, “leaders” in the population [29], individual heterogeneity [30], or flowing environments [31]. We refer the interested reader to the review by Berdahl *et al.* for a more detailed summary [9]. Whale-specific models have been presented to investigate, for example, the migration of humpback whales [32], and the migration and foraging behaviour of blue whales [33]. However, while certain realistic aspects of whale migration have been included in these models, the importance of collective navigation in such models remains to be explored, despite its conjectured importance [5, 13, 14]. In particular, the presence of a spatially- and temporally-varying noise field and its interaction with the communication range of a whale population has yet to be incorporated in a mathematical model of collective navigation and migration.

The aim of this study is to assess the extent to which ambient noise can influence whale migration paths. The lack of data under controlled conditions (as noted “baleen whales are reticent laboratory subjects” [5]) and the infeasibility of reverting from the current soundscape to a pristine soundscape motivates our computational modelling approach. Here we explore how anthropogenic activity may negatively impinge on the navigating ability of whales. Specifically, we generate synthetic migration paths while systematically addressing three possible consequences of higher noise: reduced communication space (‘solitude’), reduced goal-targeting information (‘confusion’), and the triggering of explicit reorientation responses (‘avoidance’). These plausible repercussions of increased noise levels are built into an agent-based mathematical model for whale movement that incorporates multiple layers of environmental data.

## Results

Our mathematical model is designed to simulate a virtual population of baleen whales as they migrate across the North Sea; for example, a population returning from a feeding area. While the model does not explicitly describe a particular whale species, we have parameterised the model as much as possible from data of minke whales (*Balaenoptera acutorostrata*); we note that the model could be parameterised via other whale species and other migration routes, given suitable data. The agent-based model builds on a collective navigation model introduced in [12], which assumes that the confidence in the target direction of each whale combines local inherent information, e.g. navigation cues, (Fig. 1(a)) and the collective information gained by co-aligning movement paths with other whales in communication range (Fig. 1(c)).

**Fig. 1.**
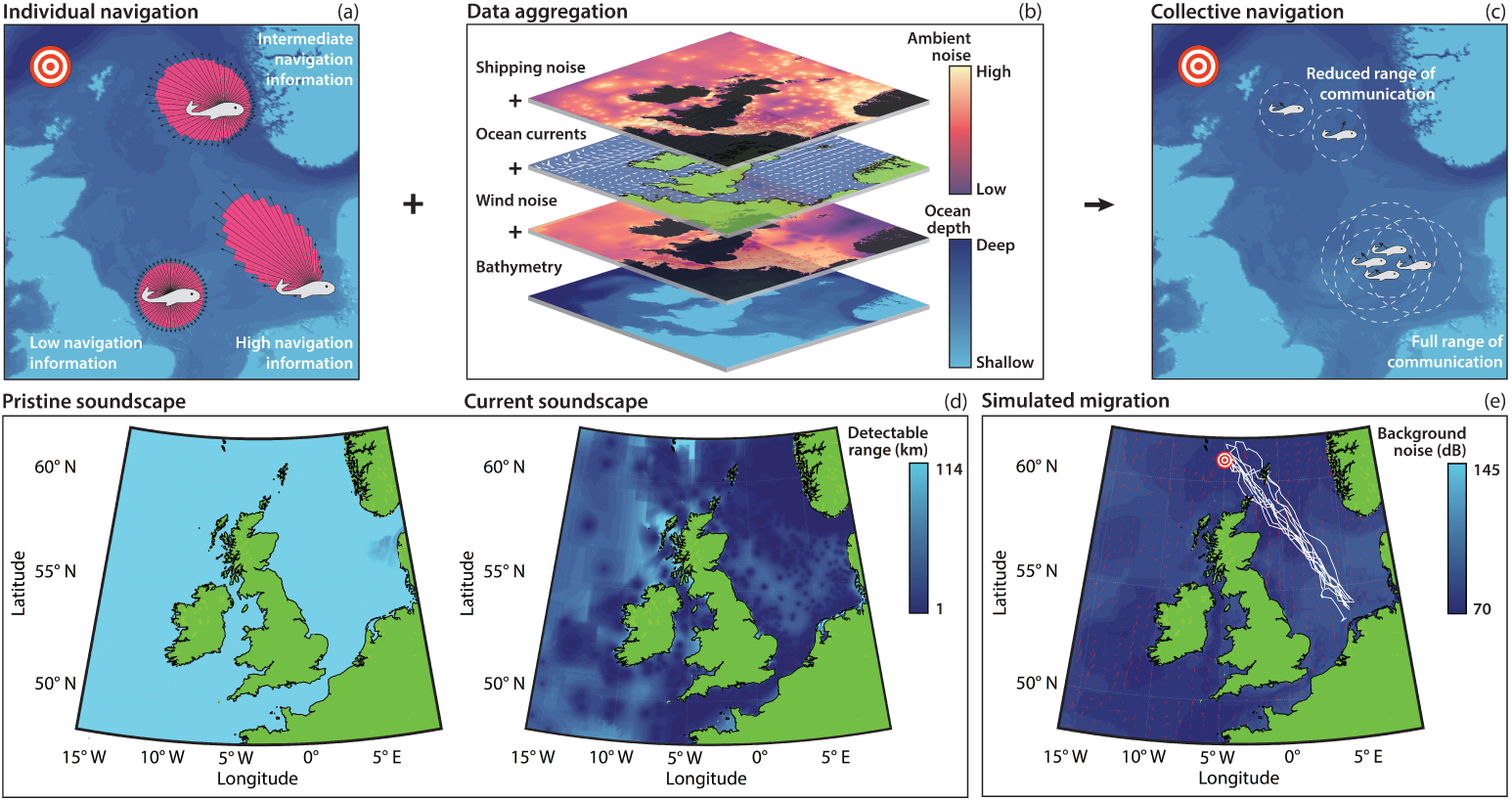
Data-driven agent-based modelling framework. (a)-(c) Schematic of the data-driven agent-based model of whale migration. (a) Individual navigation relies on inherent navigation that may be impacted by local factors. (b) Data sources incorporated in the model. (c) Collective navigation relies on the detection of signals from other whales, which is influenced by the local ambient noise. (d) The detectable range of whale calls for our model in a pristine soundscape (wind noise only) and the current soundscape (wind and shipping noise). (e) Representative, randomly-selected, trajectories from the model of collective navigation in a pristine soundscape.

In the model, the communication range is principally dictated by the ocean ambient noise (Fig. 1(d)); other factors include the sound transmission decay and source level. We decompose the ambient noise into surface wind and shipping noise levels, which allow us to consider two forms of ocean soundscape: the *pristine* soundscape (wind noise only) and the *current* soundscape (wind and shipping noise) [34]. These noise layers represent two of the four data sources needed for model implementation (Fig. 1(b)); the other data are ocean currents [35] and bathymetry [36]. A detailed explanation of the model and the parameters used, and a sensitivity analysis can be found in the Methods section and Supporting Information, which includes an Overview, Design Concepts and Details protocol document [37, 38]. The model relies on noise response mechanisms that are, by necessity, speculative due to our current understanding of baleen whales. The conclusions arising from our model should therefore be interpreted with this in mind. However, the clear qualitative differences in migration patterns that emerge from our model imply that it may be possible to identify the dominant type of response to noise from observational data in future.

### Migration in the pristine soundscape

The pristine soundscape serves as the benchmark, representing navigation in a pre-industrial ocean and offering optimal conditions for navigation: communication is close to maximum across the migration route (Fig. 1(d)). Representative migration trajectories show broadly straight line movements towards the target (Fig. 1(e)). At a population level, trajectories are constrained to a relatively tight corridor (Fig. 2(a)), implying that a high degree of cohesion is maintained throughout migration, despite the lack of an explicit “attraction” mechanism. However, this does not imply physical proximity: average pairwise distance is *∼* 100 km and only *∼* 5% of the group lie within 5 km of each other, so visual sightings of pairs or groups would form relatively rare events. The mean distance to the destination decreases linearly (Fig. 2(d)), with the majority of whales arriving within a few days of each other (Fig. 2(c)). The tail is attributed to a few “straggling” outliers, such as those either positioned at the group edge or adopting routes that require navigation about obstacles such as islands. The number of detected whale calls indicates the level of communication. This is initially high while whales are co-located at the feeding ground, but drops with migration as the group becomes dispersed. Crucially, the number of calls remains high enough to benefit navigation. We do not speculate on the rate of calls for individual whales. Rather, we assume that a whale calls more often than it undergoes reorientation. Note that during the final stages, the bathymetry helps to funnel the population between land-masses, illustrating a geophysical influence on group structure and navigation; here, land avoidance behaviour can dominate by orienting individuals away from excessively shallow waters to prevent beaching (SI Fig. 3).

**Fig. 2.**
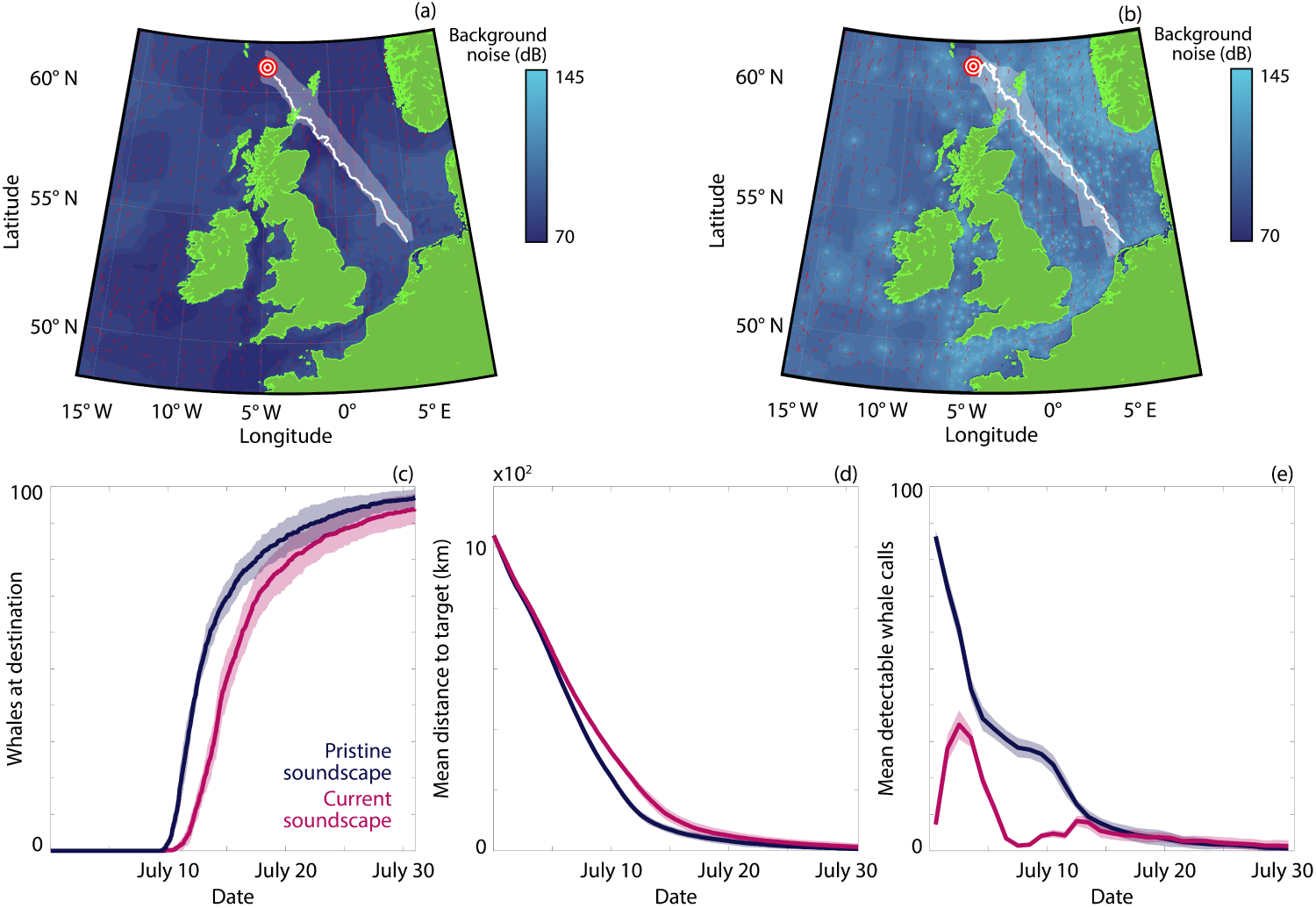
Comparison of navigation in the pristine and current soundscapes. (a) Median and spread of whale trajectories in the pristine soundscape. (b) Median and spread of whale trajectories in the current soundscape. (c) The number of whales that have arrived at the target destination. (d) The mean distance of the population from the target. (e) The mean number of detectable whale calls (averaged daily). Results are presented for the pristine (dark blue) and current (magenta) sound-scapes. The lines and ribbons correspond to the mean *±* one standard deviation over 10 simulations.

**Fig. 3.**
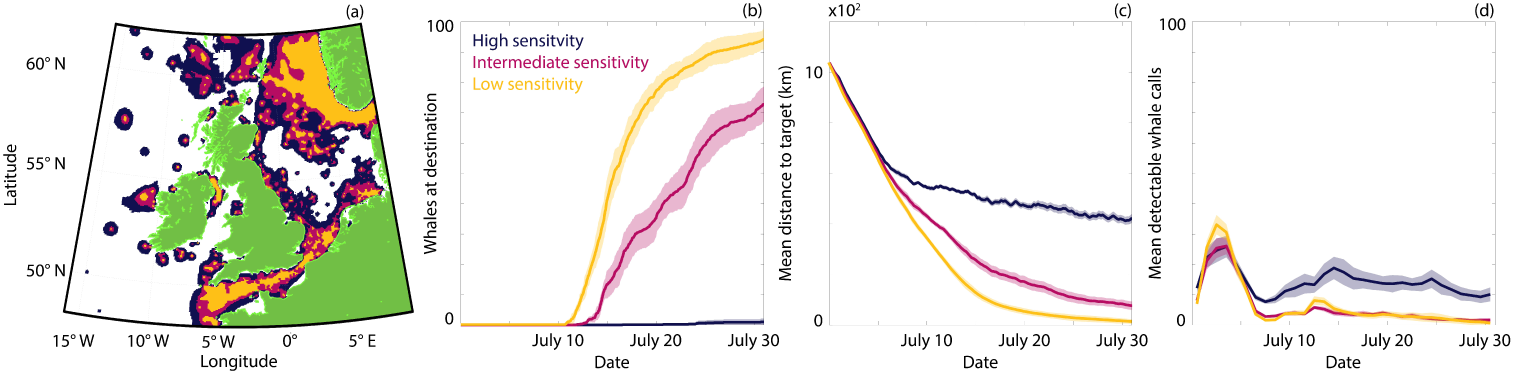
Comparison of navigation with different sensitivities to noise that induce an avoidance response. (a) Regions of the ocean that induce a noise avoidance response under low (orange), intermediate (orange+red) and high (orange+red+dark blue) sensitivities. (b) The number of whales that have arrived at the target destination. (c) The mean distance of the population from the target. (d) The mean number of detectable whale calls (averaged daily). Results are presented for high (dark blue), intermediate (magenta) and low (orange) sensitivities. The lines and ribbons correspond to the mean *±* one standard deviation over 10 simulations.

### Reduced communication range delays arrival

We now consider the current soundscape, which includes shipping noise (Fig. 2(b)) [34]. The increase in ambient noise slows migration, introducing a delay via a 3-4 day shift in the distribution of arrival times (Fig. 2(c)). This effectively represents an additional *∼* 20% in travel time. Longer migration primarily results from reduced communication space, as the detectable range drops by an order of magnitude or so across the migration (Fig. 1(d)). This manifests in a dramatic decrease in the number of detectable calls (Fig. 2(e)), when compared against the pristine soundscape. This is particular apparent from the outset, where initial proximity to shipping lanes leads to considerably reduced communication. Calling recovers as the population moves into quieter seas. The funneling between landmasses towards the end of the migration proves particularly beneficial here, bringing individuals within communication range despite the noise-induced masking. Overall, though, throughout migration the communication range is greatly diminished and the advantages of collective navigation are lost, with individuals more heavily reliant on inherent information such as memory or cue detection.

### Noise avoidance can block migration routes

Explicit noise avoidance behaviour is included in Fig. 2, but is only triggered at higher exposure levels. Entirely eliminating this avoidance only marginally improves navigation (SI Fig. 1), indicating that the impaired navigation observed in Fig. 2 primarily stems from communication masking. Clear noise avoidance responses are well documented for whales exposed to nearby loud noise sources and are typically based on direct visual observations [23]. As such, uncertainty exists regarding the level of noise at which such a response occurs, with more subtle path deviations difficult to assess visually. To explore the impact of explicit noise avoidance we lower the intensity threshold at which this behaviour is triggered (Fig. 3) (changing from *∼*1 km to *∼*8 km from a large ship). Navigation efficiency is significantly reduced, with a slower approach to the target and delayed arrival times are observed (Fig. 3(b-c)). Notably, significant differences only emerge around halfway through the journey. To understand this we chart the regions where noise avoidance is triggered for different sensitivities (Fig. 3(a)). The first portion of the migration path remains relatively clear for all sensitivities, yet later stage migration becomes more convoluted due to higher ambient noise. Consequently, routes that do not require noise avoidance become restricted to certain channels, which can be closed off as the sensitivity increases. This, in turn, leads to failures in migration (Fig. 3(b-d)), as individuals become trapped behind a wall of noise. We further explore this phenomenon by systematically varying the parameters in the noise avoidance mechanism (Supporting Information).

### Noise-induced reductions in inherent information

Higher ambient noise may also reduce the level of inherent information available. This may occur either directly, through obscuring sound sources that serve as navigation cues; or indirectly, through poor processing of information due to noise-induced confusion or stress. Notably, we observe significant delays in arrival time when noise-induced information loss is included, becoming severe if the loss is high (Fig. 4). Crucially, this ineffective navigation does not result from closed routes but from less-directed movement that places whales at greater susceptibility to dispersing effects, including currents. This is particularly prominent across the second half of the migration, where the higher noise within this region results in greater spread and significant deviation of the median trajectory (Fig. 4(a)). Previous studies indicate that poor information zones can be compensated for through collective navigation [12], via a communication relay between information-rich and information-poor regions. Here, this compensation is unavailable as noise concurrently reduces group communication. Hence, in the model, an increase in ambient noise can have a compounding and negative impact on navigation efficiency due to simultaneously affecting the different elements that aid navigation.

**Fig. 4.**
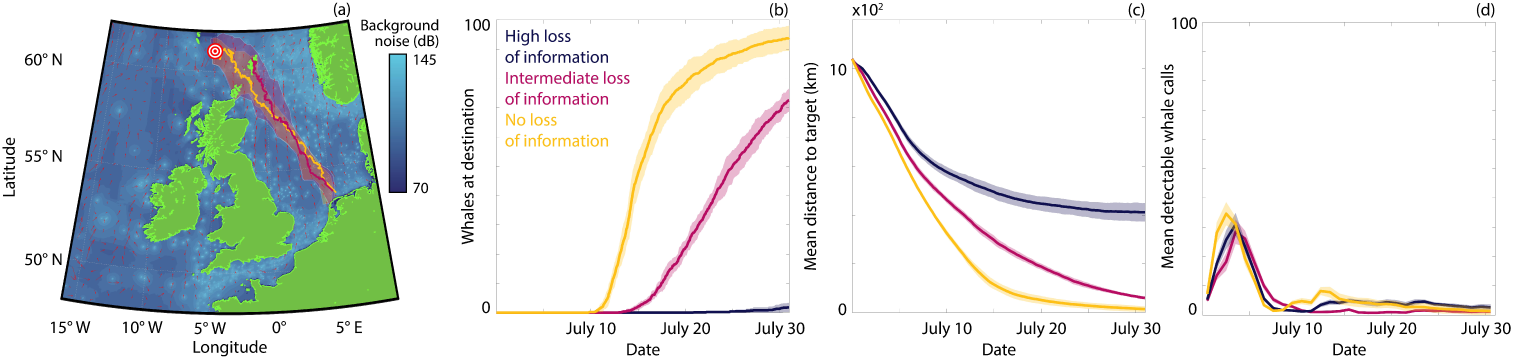
Comparison of different levels of loss of inherent information due to noise pol-lution. (a) Median and spread of whale trajectories. (b) The number of whales that have arrived at the target destination. (c) The mean distance of the population from the target. (d) The mean number of detectable whale calls (averaged daily). Results are presented for high (dark blue), inter-mediate (magenta) and no (orange) loss of inherent information. The lines and ribbons correspond to the mean *±* one standard deviation over 10 simulations.

### Perturbing the current soundscape

Building on the insights of the above analysis, the predictive potential of the model is demonstrated through perturbing the current soundscape (Fig. 5). Specifically, we (i) construct synthetic noise maps with shipping noise that originates from virtual vessels (Fig. 5(a)), and; (ii) consider the localised impact of a large scale offshore construction process in an otherwise pristine soundscape (Supporting Information). For the synthetic equivalent to the current soundscape we set parameters for the routes, numbers and source levels of virtual vessels to near recreate the migration behaviour within the current soundscape profile (Fig. 2). From this baseline we consider the impact of a future 50% increase in traffic (Fig. 5, dark blue). The resulting rise in ambient noise hinders migration through triggering the noise avoidance that traps a subset of the population behind high noise regimes. As a potential mitigation we explore the extent to which introducing slowdown zones can offset the impaired migration; trials indicate that slower speeds can reduce noise levels from certain vessels upwards of 10 dB [39]. Introducing a slowdown can partially recover migration efficiency (Fig. 5, magenta), despite the elevated vessel numbers. To simulate the impact of a construction project, we place a single noise source at a fixed location and consider different amounts of construction activity (e.g. pile driving) per day. Increasing construction activity leads to an increasing perturbation to the migration path (Fig. 6(a)) and day-to-day activity triggers oscillations in the remaining migration distance (Fig. 6(c)). The latter results from a daily triggering of noise avoidance behaviour that operates until the whales have moved sufficiently far away from the source to allow normal migration. The total amount of construction activity has a more pronounced impact on migration than the scheduling of the activity (Supporting Information).

**Fig. 5.**
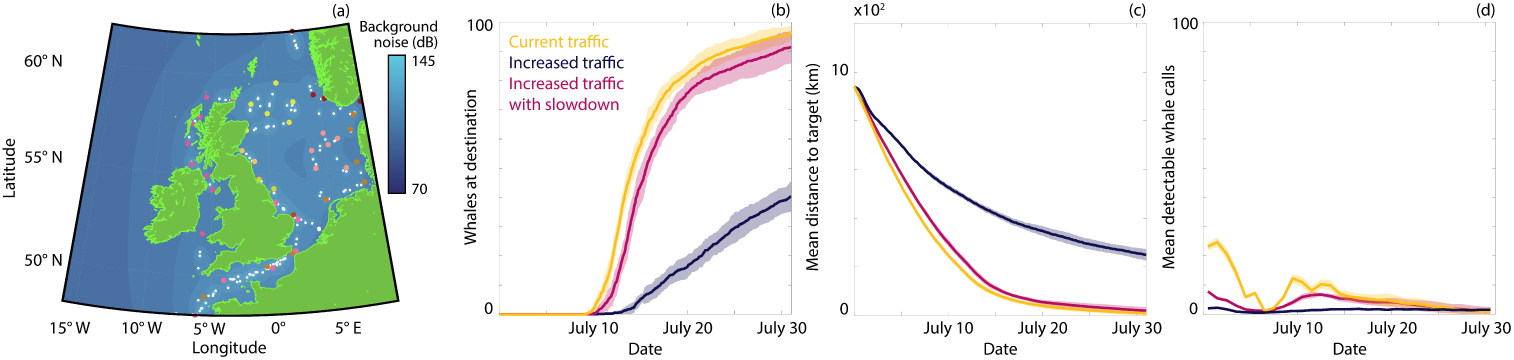
Comparison of navigation under different levels of shipping activity. (a) Represen-tative noise map for shipping traffic. Start points, end points and waypoints for individual routes are shown in unique colours; ship locations are white. (b) The number of whales that have arrived at the target destination. (c) The mean distance of the population from the target. (d) The mean number of detectable whale calls (averaged daily). Results are presented for current shipping traffic (orange), increased traffic (dark blue) and increased traffic alongside noise mitigation measures (magenta). The lines and ribbons correspond to the mean *±* one standard deviation over 10 simulations.

**Fig. 6.**
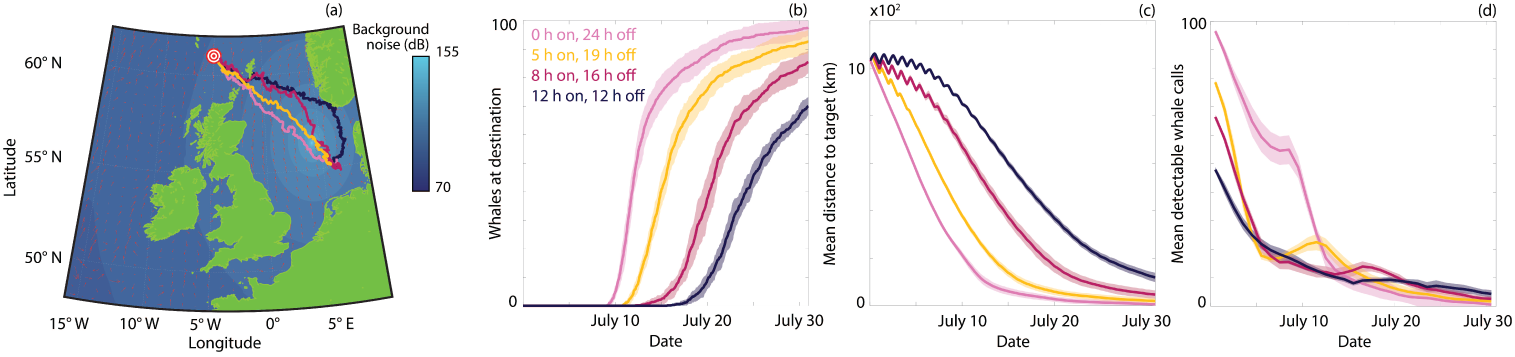
Comparison of navigation under different levels of construction activity in an otherwise pristine soundscape. (a) Noise map during active construction at 56.35 *◦*N, 4.25 *◦*E with median trajectories. (b) The number of whales that have arrived at the target destination. (c) The mean distance of the population from the target. (d) The mean number of detectable whale calls (averaged daily). Results are presented for 0 hours (pink), 5 hours (orange), 8 hours (magenta) or 12 hours (dark blue) of construction activity per day. The lines and ribbons correspond to the mean *±* one standard deviation over 10 simulations.

## Discussion

We have developed a model to explore the impact of ocean noise on whale migration routes, where higher ambient noise (i) reduces whale communication space, (ii) generates an avoidance response when sufficiently loud, and (iii) lowers inherent navigation information. Each mechanism can lengthen the journey time, and certain scenarios may even lead to failed migration. As such, the energetic cost of migrating in the current ocean soundscape is expected to be higher than in a pristine soundscape. Notably, though, each mechanism has a subtly different impact: it is not simply three different forms of slower migration. Rather, diminished communication space leads to greater solitude and slower migration as collective navigation benefits are eroded. Under a loss of information there is increased confusion, leading to off-course drifting and greater susceptibility to ocean currents. Finally, loud noises lead to a strong noise avoidance response and routes that become blocked, and hence migration may fail. Whether these distinct forms of trajectory perturbation predicted by our virtual whale model are also observed within tracked whales would be of key interest. The conclusions drawn from the model predictions are contingent on the relevance and accuracy of the mechanisms in the model. It is likely that baleen whales exhibit more sophisticated behaviour than that encoded into the model, including learning-type responses and adaptation to the evolving environmental conditions. However, in the absence of detailed understanding we sought to impose relatively simple mechanisms and explore how these mechanisms may manifest in qualitatively different migration behaviour.

Reduced communication spaces lengthen journey times through lower collective navigation. Previous theoretical studies [11, 12] indicate that co-alignment of migration paths provides a “many wrongs” [10] benefit by reducing inherent uncertainty, but only above a critical number of detectable neighbours [11, 12]. A reduced communication space can therefore eliminate this benefit, leading to a more convoluted path. We have not (explicitly) included noise compensation behaviour, such as the Lombard effect. This effect has been observed in whale populations, where call intensities are increased at higher ambient noise levels [24, 25]. It would be possible to include this mechanism, yet it is known to provide only provide partial compensation [25], and we would expect qualitatively similar results from the model.

Allowing ambient noise to reduce inherent information also negatively impacts on migration times. The guidance cues used by whales during their navigation are largely unknown, but listening for characteristic sound sources is certainly plausible [3, 7]: surf may allow detection of coastlines [6], while rifting of icebergs creates noise sources estimated at *∼* 245 dB (re 1 m) [40], detectable thousands of kilometres away for the low frequency bands of baleen whale acoustics. Higher ambient noise therefore may reduce the detectability of such sources, decreasing the efficacy of target-directed motion and increasing the susceptibility of whales to ocean currents. It is also feasible that louder ambient noise reduces inherent information from other sources, such as geomagnetic field information [3], through a noise-induced confusion that lessens processing ability.

A more direct and acute response included here is noise avoidance [22, 23], where noise levels above a threshold induce movement away from the source. This can impact significantly beyond introducing deviations into migration paths. At the extreme end, migration routes may become blocked if whales are “trapped” behind a wall of noise. Noise avoidance is modelled as a ‘negative phonotaxis’ response: directed away from loud noise sources. Consequently, this acts to concentrate the population within lower noise regions, where avoidance responses are not triggered. Evidence for a noise-induced spatial redistribution of whale populations can be found in [41], following analyses into the tracks derived from detected minke whale calls before, during, and after periods of naval activity [41]. Statistical modelling further supports the hypothesis that individuals were specifically moving away from sonar-producing ships [22]. Negative phonotaxis responses could conceivably be achieved through a comparison of current noise levels with prior noise levels.

We have currently assumed a constant swimming speed. Noise avoidance, however, may also manifest in (substantial) increases to the movement speed as the individual escapes [23]. Higher speeds carry significant extra costs: energy expenditure during migration stems from both metabolism and generating propulsion. For marine animals the latter rapidly increases with speed due to the increased drag [42, 43]. Optimal migration speeds that minimise energy expenditure can be calculated for different cetacean species [44, 45]. Therefore, extending the model to include noise-modulated speed and tracking energy expenditure will allow further insight into how noise impacts on whale fitness.

More severe reactions to noise are possible beyond those included here. Numerous studies imply a causal relationship between extreme noise events and cetacean mass strandings, with close spatiotemporal correlations and biopsies indicating noise-induced physiological damage [19]. A whole spectrum of reactions is therefore plausible, from subtle to severe. Highly elevated stress could result in disorientation, potentially impacting the land avoidance mechanism that prevents beaching. Exposure to loud noises may result in transient or permanent hearing impairment, debilitating whales beyond the time of exposure [19]. These factors could be included through an additional variable, describing a whale’s current physiological state. Population heterogeneity could also be included via an age variable, that impacts both on hearing sensitivity (e.g. age-related hearing loss in older cetaceans, e.g. [46]) and navigating ability (e.g. less experienced juvenile members [3]). Incorporating these and other forms of heterogeneity will allow us to explore whether ambient noise disproportionately impacts on different members of the group.

Sound information has been condensed here into the intensity level, allowing us to use a simple transmission loss model [20] in which calls are heard only when the received intensity is not sufficiently below the ambient noise. A more sophisticated model could account for noises covering different frequency bands and species-specific sensitivities. However, while available for certain other cetaceans, audiograms have not yet been obtained for baleen whales [20] and the simpler intensity-based model would appear a reasonable compromise at present.

A number of other studies have also used agent-based modelling to explore the impact of noise on whale communication space or navigation. For example, Cholewiak *et al.* [26] consider a model for mobile whales that swim within a region subject to shipping traffic, exploring the change in communication space for different calls across a variety of baleen populations and according to the different forms of shipping. Guarini and Coston-Guarini [32] explore the influence of bathymetry on humpback whale migrations. The work here extends the preliminary study performed in [12], and merges aspects of navigation with impacts from communication masking, while also accounting for the effects of ocean currents and bathymetry. Ocean currents can reach orders of magnitude commensurate with migration speeds and have been accounted here as a passive advection. This assumes that whales do not specifically adapt their movement with respect to an ocean current – we are not aware of any studies that suggest whales orient according to the current (rheotaxis). Nevertheless, aligning with currents is common within marine animals [47], and this could be included in future. Land avoidance was primarily included in our model to prevent beaching. However, it was also found to confer navigating benefits by funneling whales between landmasses, improving cohesion in the process.

Our case study adopted the North Sea region due to its high level of human activity, the availability of data (noise maps [34], bathymetry [36], ocean currents [35]) and the presence of various cetaceans [48], including a seasonal aggregation of minke whales [49]. However, the modular nature of the model allows it to be adapted to other case studies: for example, noise maps are also available for Australian coastal waters [50], where humpback whales migrate along the eastern coast to breeding grounds in the Great Barrier Reef [51]. The model can also be used as a prediction tool, illustrated here by considering beneficial (lowered shipping speeds) and detrimental (increased traffic, introducing offshore construction projects) perturbations to the soundscape. It would also be possible to extend the model to consider a spectrum of potential climate change impacts, which could include altered ocean currents, changes in sound transmission due to ocean acidification or spatiotemporal changes to a target food resource. Simulating a sequence of migrations over multiple years across an age-structured whale population would allow exploration into the potential resilience of whale populations in the light of such changes.

## Materials and methods

### Study site

Our study considers the movement of a hypothetical baleen whale population along a (predominantly) south to north route across the North Sea, from a region north of the Netherlands toward the Atlantic Ocean between Scotland and the Faroe Islands. Our focus on this region is motivated by the considerable human activity in these waters, with significant levels of shipping, exploration and construction. As a by-product to this activity, there is an availability of fine-scale data (ocean noise [34], bathymetry [36] and ocean currents [35]) that form key inputs into the model. To allow fixing of certain parameters of the model (e.g. migration speeds and call source levels) we consider data for minke whales (*Balaenoptera acutorostrata*), which is the most populous species of baleen whale within the North Sea [52, 53]. The starting location coincides with previous observations of a significant seasonal aggregation near the Dogger Bank [49] region that peaks in May. However, we stress that we are not specifically modelling this species: the overall aim is to understand the potential impact of environmental factors, principally noise, on navigation. However, the model framework is flexible and can be tailored to explore specific whale migrations, given appropriate data.

### Model

The basis for the model presented here is the collective navigation model in [12]. Full details can be found in the original manuscript; however, we briefly summarise the model here. Each virtual whale is tracked according to its position (x) and swimming orientation (*θ*), and moves according to

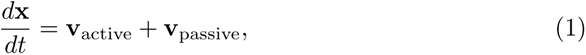

where **v**_active_ represents the contribution to its velocity from active swimming and **v**_passive_ is the contribution from ocean currents. Each individual undergoes active motion according to a velocity-jump random walk [54]. Individuals swim with a fixed heading (*θ*) and speed (*s*) for an exponentially-distributed length of time before selecting a new heading, and repeating the process. The heading selection process encodes collective navigation as the selected heading reflects the inherent knowledge of a target in combination with the observed heading of other nearby individuals [12]. Crucially, navigational uncertainty is captured by the spread in the set of observed headings. This previously-proposed model effectively demonstrates the benefit of collective navigation, particularly in regions of poor navigation information. However, certain model features necessary for describing long-distance whale migration are not present in the previous model. We extend the model to incorporate the influence of ocean currents, dynamic noise pollution and signal detection, noise avoidance, and land avoidance. In all simulations we consider 100 whales. All simulations are performed in Matlab R2020b. The code used to perform the model simulations can be found at https://github.com/DrStuartJohnston/whale-migration-model.

### Ocean currents

Ocean currents can benefit (or hinder) migration simply via passive transport according to whether the target direction is aligned (or opposed) with the dominant current direction. In certain regions, current velocities can reach speeds comparable to average migration speeds, and hence form a non-negligible contribution to motion that is incorporated through the passive transport contribution of (1). The ocean current data used here is obtained from the HYCOM model [35], which provides day-to-day currents at a spatial resolution of 0.04*^◦^* latitude and 0.08*^◦^* longitude. We only use the surface depth layer of this data: while baleen whales do make occasional deeper dives, we presume that while migrating they remain close to the surface, alternating between swimming and surfacing to breathe.

### Ocean noise

Ocean noise is a product of both natural processes, such as wind and rain, and anthropogenic activity, such as shipping and resource exploration, each varying spatially and temporally [17]. Shipping activity is concentrated along commercial shipping lanes, while rain occurs in transient bands. Heightened awareness of the importance of the ocean soundscape has led to increased acoustic monitoring [55] and the generation and validation of ‘soundmaps’ [34, 56]. These maps require significant computational over-head and, consequently, we utilise the data from these previously-published studies rather than explicitly modelling spatial noise distributions. Specifically, we use data generated in [34], which provides validated estimates for the noise levels across the North Sea that arise from shipping traffic and wind at the ocean surface. Separating this data according to the distinct noise sources subsequently allows us to consider both pristine (wind noise only) and current (both shipping and wind noise) sound-scapes. This data also informs the parameterisation of synthetic noise maps (i.e. noise maps based on simulated traffic) based on a simplified sound transmission model. Specifically, the received level RL (dB) at a distance *r* (m) from a sound with source level SL (dB), can be modelled by

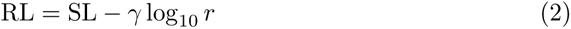

where *γ* log_10_ *r* describes the transmission loss [20]. The coefficient *γ* is bounded below by 10, corresponding to shallow water in which the spreading is effectively cylindrical, and above by 20, for deep water in which sound propagates in all directions (spherical). Here we use *γ* = 17.8 [57]. This is a considerable simplification; however, this makes it feasible to construct and analyse multiple synthetic noise maps, which would not be the case for detailed sound transmission models (which require the solution of partial differential equations across the ocean, accounting for accurate bathymetric data) [34]. It is plausible that baleen whales are more sensitive to specific noise bands, particularly lower frequency bands [58], and this remains a model extension of interest, if a balance can be found between fidelity of sound transmission physics and computational complexity.

### Communication range

We explicitly model potential communication masking [20], in which the call produced by a whale may be masked by the ambient noise. Specifically, we assume that a whale emits a call at 178 dB (re 1 m), consistent with median estimates of minke whale pulse trains [59], and that the transmission of this sound follows (2). This call can be detected by another individual if the signal-to-noise ratio (SNR) of the signal and the ambient noise at the location is above a theshold value

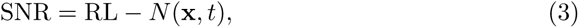

where *N* (x*, t*) is the ambient noise at location x and time *t*. As above, this is a considerable simplification of the complex nature of real world whale communication, where calling, transmission and detection will be frequency and orientation dependent. Plausible SNR are not restricted to positive values; for example, negative values are observed for human conversation in noisy environments [60].

### Noise avoidance

Loud noise sources have been observed to induce escape or avoidance behaviour [61]. For example, see [22, 23] with specific reference to minke whale noise avoidance responses to sonar. To include these responses into the model, we impose a mechanism where individuals reorient and move away from regions of high noise (negative phonotaxis). The strength of this behaviour is given by a sigmoidal function such that (i) below a threshold noise level there is essentially zero noise avoidance, (ii) for intermediate noise levels individuals balance noise avoidance against navigation, (iii) above a certain threshold, navigation behaviour is neglected and motion is dictated solely via noise avoidance. This latter could be viewed as essentially a stressed or panic response, where the noise overrides other factors [19]. The detailed functional form and sensitivity analysis of the noise avoidance mechanism can be found in the Supporting Information.

### Land avoidance

We assume healthy whales, under normal noise levels, avoid shallow water to minimise risk of beaching. To include this behaviour, we implement a similar sigmoidal approach to land avoidance as for noise avoidance. Specifically, we use bathymetric data [36] to obtain estimates of the water depth at any given spatial coordinate; whales likely estimate distance to the ocean floor through specific downward-directed calls [7], or to the shore through listening for surf [6]. For deep water, land avoidance is essentially not considered. If the individual is in sufficiently shallow water, however, navigation is neglected and the individual prioritises motion in the direction of deepest water (we call this “bathotaxis”). At intermediate depths, there is a weighting between navigation and land avoidance.

### Data sources

As noted above, implementation of the model requires synthesis of multiple datasets: ocean current velocity data from HYCOM Global (GLBv0.08) [35], bathymetry data from EMODnet Digital Bathymetry [36], coastline data from the Global Self-consistent, Hierarchical, High-resolution, Geography Database [62], and wind-derived noise from [34]. Shipping-derived noise data is either obtained from the study [34] or from our synthetic noise model. The synthetic noise model allows incorporation of both fixed (e.g. drilling and exploration) and mobile (e.g. ships) noise sources of varying intensity, therefore permitting investigation into the influence of different shipping lanes, levels and types of traffic, or the introduction of new construction projects. We use linear interpolation between data points to evaluate each of the datasets at the location of an individual in the model. All simulations are performed in Matlab R2020b. The code used to perform the model simulations can be found at https://github.com/DrStuartJohnston/whale-migration-model.

## Acknowledgements

We thank Adrian Farcas for supplying and assistance with the North Sea ambient noise data used to generate sound maps.

## Supplementary information

The Supporting Information consists of two documents. The first provides a more detailed description about the modelling framework and contains additional results, including a sensitivity analysis. The second is a Overview, Design Concepts and Details protocol document, which presents the details of the agent-based model in a standardised format.

## Declarations

### Ethics approval

Not applicable.

### Consent to participate

Not applicable.

### Consent for publication

All authors consent to the publication of the manuscript.

### Availability of data and materials

The data used as inputs for the simulations can be found at https://github.com/DrStuartJohnston/whale-migration-model.

### Code availability

The code used to perform the model simulations can be found at https://github.com/DrStuartJohnston/whale-migration-model.

### Author contributions

STJ: conceptualization, formal analysis, investigation, methodology, project administration, resources, software, validation, visualization, writing -original draft, writing — review and editing; KJP: conceptualization, formal analysis, investigation, methodology, validation, writing - original draft, writing — review and editing.

## Supplementary information

### Model and parameter details

For all simulations we consider 100 whales. While we do not model a specific migration, parameters are fixed according to data for minke whales; this species is relatively abundant within the North Sea [6] and a significant subpopulation has been observed to congregate at feeding grounds at the southern end of the North Sea during spring/summer [3]. In our model the agent whales are initially distributed at random within a prespecified region, broadly compatible with the localisation of the congregation noted above and ensuring that initially the population is within communication range. For all results except Figure 5, this is the region defined by 53.5*^◦^* N to 54.5*^◦^* N and 4.5*^◦^* E to 5.5*^◦^* E. For Figure 5, this region is defined by 55*^◦^* N to 56*^◦^* N and 4.5*^◦^* E to 5.5*^◦^* E. The target destination of the whale population in all cases is 61*^◦^* N, 5*^◦^* W. Whales are considered to have arrived if they are within 50 km of the target destination.

For all simulations we assume a constant speed of 6 km/h, within the estimated range of migration speeds for minke whales [8]. We assume a reorientation rate of, on average, once per hour. The time between reorientation events is exponentially distributed. Between reorientation events whales move with a constant velocity and the new heading selected during the reorientation process follows the procedure described in the model by Johnston and Painter [10]. Consistent with that model we select *α* = 0.5 and *β* = 0.5, where *α* and *β* each represent weighting parameters between the inherent and collective information when estimating the mean direction and uncertainty, respectively. This choice reflects the results from an exploration in [10], where an even weighting between inherent and collective information is determined to give rise to near-optimal navigation. We select a background level of inherent information with *κ* = 1; again consistent with the study in [10], this value ensures that the level of inherent navigation information can capably guide whales towards the target in the absence of collective navigation, but that collective behaviour can provide a significant boost to journey times.

Each simulation is conducted across 744 (model) hours, i.e. 31 (model) days. Note that we assume that whales do not break their migration (e.g. to rest, feed or sleep). We set the migration to begin on July 1 2010, and use the corresponding ocean current data from the HYCOM Global (GLBv0.08) model [2] with a time spacing of one day.

We assume that the source level (SL) of the whale call is 178 dB (re 1 m), consistent with previous estimates [13]. We choose a signal to noise ratio (SNR) for detection of whale calls among background noise of SNR = *−*5 dB. The minimum received level for a call to be detected is chosen to be 88 dB (re 1 m). Under our sound transmission model, this provides a (pristine) communication range of *∼*114 km, a value chosen for its consistency with representative values of potential communication space in other studies of minke whales [7]. Note that there is an absence of audiogram data/hearing sensitivity for baleen whales [4], including minke whales, leaving uncertainty regarding their actual communication range.

To describe noise avoidance behaviour we implement a negative phonotaxis response (i.e. motion in the direction of decreasing noise). This response is implemented in a weighted manner, where at low noise levels there is essentially no response, and at high noise levels, essentially all motion is determined by the noise avoidance response. Specifically, we calculate the proportion of motion that is driven by noise avoidance *w_na_*(*N* (x*, t*)), where the *N* (x*, t*) is the noise level at location x and time *t*. The remaining proportion of motion (i.e. 1 *− w_na_*(*N* (x*, t*))) is motion corresponding to regular migration behaviour. We calculate the weighting via

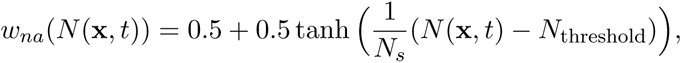

where *N*_threshold_ is a threshold parameter that represents the noise level at which there is an equal weighting between noise avoidance and migration and *N_s_* is the transition rate parameter that dictates the rate of transition between regular and noise avoidance behaviour. Here *N*_threshold_ = 120 dB (re 1 m) for Figures 2 and 4, *N*_threshold_ = 110 dB (re 1 m) (dark blue), 115 dB (re 1 m) (magenta), 120 dB (re 1 m) (orange) for Figure 3 and *N*_threshold_ = 115 dB (re 1 m) for Figure 5. We demonstrate the efficacy of the noise avoidance response in Fig. 2, where we place an (artificial) extreme noise source in the centre of the North Sea. We observe a clear noise avoidance response, where trajectories indicate that the whales skirt around the edge of the region of extreme noise, and then recommence normal migration behaviour.

We consider a similar approach for land avoidance, assuming that excessively shallow water will trigger a response in which motion is in the direction of greatest water depth (bathotaxis). Specifically, we define a weighting *w_la_*(*d*(x)) that represents the proportion of motion that is in the direction of greatest water depth, given the depth at the current location *d*(x). Similar to the noise avoidance response, the remaining proportion of motion (i.e. 1 *− w_la_*(*d*(x))) is motion corresponding to regular migration behaviour. We calculate the weighting via

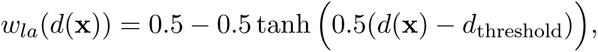

where *d*_threshold_ is a threshold depth that represents the water depth at which there is an equal weighting between land avoidance and migration. Here we assume *d*_threshold_ = 30 m. We note that in the present implementation of the model, land avoidance responses are balanced against noise avoidance responses (i.e. ensuring the total weights are no more than one); a failsafe mechanism is present, where any motion that would result in whales crossing onto land is aborted. However, the failsafe mechanism is not invoked in any of the scenarios we consider, i.e. there are no noise responses that drive whales onto land. We remark that this is an interesting avenue to explore, and would allow exploration into potential whale beaching events. We demonstrate the land avoidance response in Fig. 3, where we observe a population of whales that are able to avoid Ireland and navigate around its coast.

Generally our model assumes the level of inherent information to be constant (*κ* = 1). However, for Figure 4 we consider a potential impact in which the level of inherent information is reduced in the presence of higher noise. To implement this we choose *κ*(x*, t*) as a function of the noise:

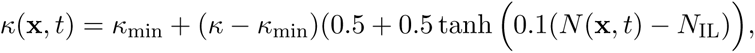

where *κ*_min_ is the minimum level of inherent information (i.e. at extreme noise levels) and *N*_IL_ is a threshold parameter that represents the noise level at which half of the inherent information (that can be lost) is lost. In Figure 4 we use *κ*_min_ = 0 and *N*_IL_ = 110 dB (re 1 m) (red), 100 dB (re 1 m) (dark blue). Note that here there is a slower transition between the two extremes (full information and no information) due to the slope parameter in the tanh function, compared to the land and noise avoidance mechanisms. This allows us to capture a range of inherent information levels across the migration.

### Model metrics

For the model output, we report up to 4 key statistics. All statistics are calculated across *N*_repeats_ = 10 identically-prepared realisations of the simulation. The first is the median trajectory of the population [1]. As the concept of a mean trajectory is not well defined in the presence of divergent paths (i.e. around obstacles), we follow Buchin *et al.* [1] and calculate the median trajectory. The median trajectory is defined by following the trajectory of an individual until it intersects with the trajectory of another individual, after which the median trajectory follows the trajectory of the other individual. The median trajectory follows this trajectory until another intersection event, after which the trajectory switches again. This approach ensures that the median trajectory always reflects a component of an actual trajectory and that the median trajectory is bound by the outermost individual trajectories [1]. We use the Matlab package “Fast Line Segment Intersection” [5] to efficiently determine intersections. We present the spread around the median trajectory by dividing space into bins and calculating the relative frequency that a whale is located in that bin. The boundary of all bins above a threshold value (0.05) is used to generate the white transparent region in the figures.

The second metric is the number of whales at the target location. This is calculated by determining the average number of whales that remain in the simulation at a time and subtracting this from the initial number of whales.

The third metric is the mean distance to the target location. This is calculated by determining the centre of the whale population in an individual simulation at each time point, and calculating the Euclidean distance between the centre of the population and the target location. This is then averaged across each simulation repeat.

The fourth metric is the average number of detected whale calls. At each time point we calculate the number of other whales each individual whale in the population can detect. To account for the rapidly varying noise level in the ocean, we calculate the average of this across the population and across individual simulation days. This metric is then averaged across each simulation repeat.

### Model parameters Shipping traffic model

To investigate potential changes in shipping traffic and the impact that this change may have on whale migration we develop a model of shipping traffic and sound transmission. We define shipping routes in our domain of interest via start and end points that are connected via a number of waypoints. These routes are selected to align with established shipping routes in the North Sea. For each route, we define the number of ships that will commence along that route over the course of 100 hours. We consider five classes of ship that contribute significantly to human activity in the oceans as per MacGillivray *et al.* [11]: bulk carriers, container ships, cruise ships, tanker ships and vehicle carriers. Under normal activity in our model, bulk carriers have a mean speed of 13 knots and a mean source level of 188 dB (re 1 m). Container ships have a mean speed of 19 knots and a mean source level of 191 dB (re 1 m). Cruise ships have a mean speed of 17 knots and a mean source level of 183 dB (re 1 m). Tanker ships have a mean speed of 14 knots and a mean source level of 187 dB (re 1 m). Vehicle carriers have a mean speed of 18 knots and a mean source level of 190 dB (re 1 m). In the slowdown scenario, each ship is assumed to reduce its speed to 11 knots with a corresponding reduced mean source level of 181 dB (re 1 m) (bulk carrier), 179 dB (re 1 m) (container ships), 175 db (re 1 m) (cruise ships) and 178 db (re 1 m) (vehicle carriers).

By combining the start points, end points, waypoints and ship properties it is straightforward to calculate the location of emitted noises corresponding to individual ships. We note that the model is extremely flexible and new routes and ships can be defined as necessary; it is also possible to include other stationary or mobile noise sources, e.g. drilling or naval sonar. We incorporate the ship locations and source levels into the logarithmic model of sound transmission discussed in the Methods of the manuscript, and hence we can calculate a noise map corresponding to the synthetically-generated shipping traffic. We assume that shipping patterns do not vary significantly over the short term and allow the pattern to repeat on the scale of 100 hours.

**Table 1:**
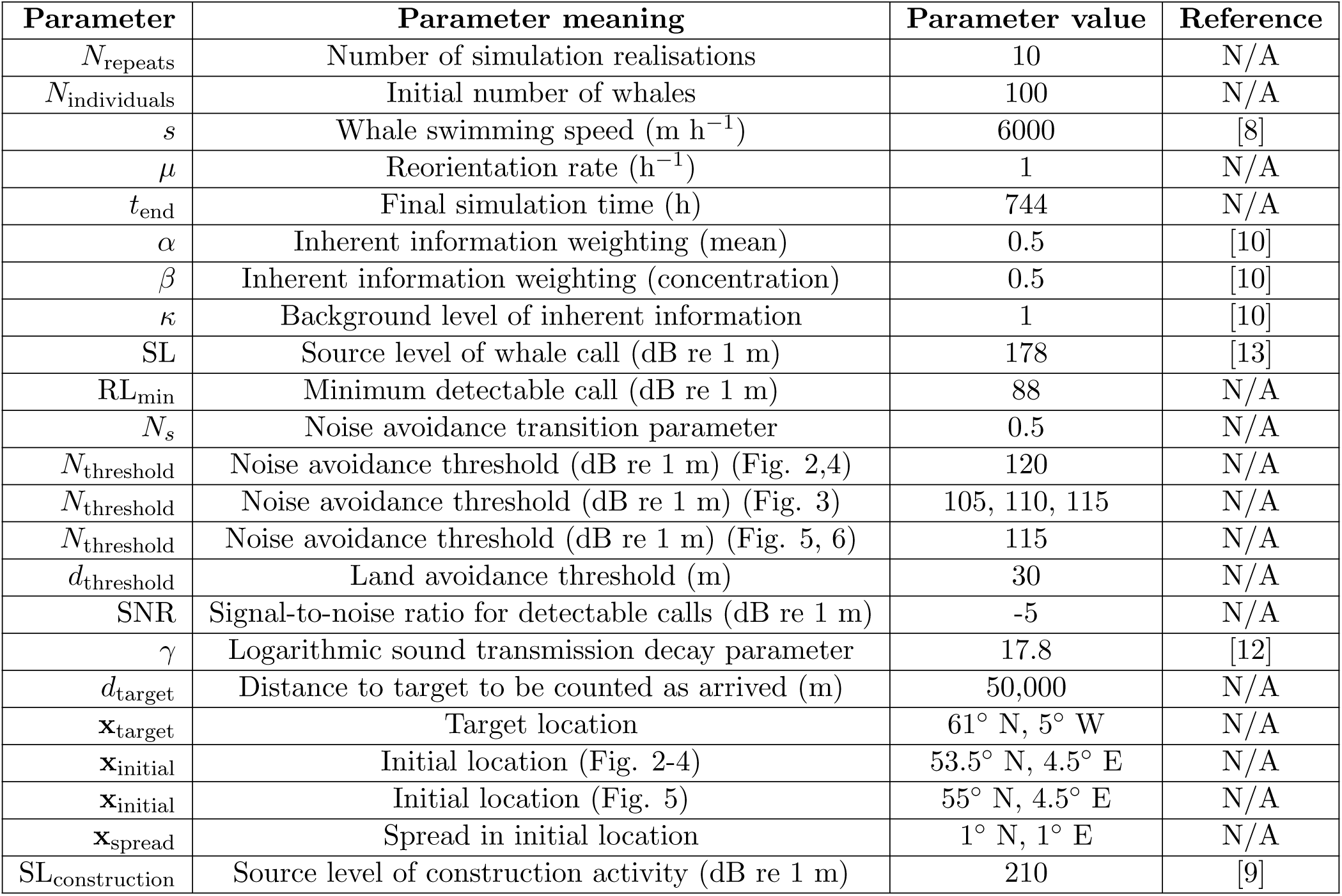
Table of model parameters.

| Parameter | Parameter meaning | Parameter value | Reference |
| --- | --- | --- | --- |
| $N_{\text{repeats}}$ | Number of simulation realisations | 10 | N/A |
| $N_{\text{individuals}}$ | Initial number of whales | 100 | N/A |
| $s$ | Whale swimming speed (m h<sup>-1</sup>) | 6000 | [8] |
| $\mu$ | Reorientation rate (h<sup>-1</sup>) | 1 | N/A |
| $t_{\text{end}}$ | Final simulation time (h) | 744 | N/A |
| $\alpha$ | Inherent information weighting (mean) | 0.5 | [10] |
| $\beta$ | Inherent information weighting (concentration) | 0.5 | [10] |
| $\kappa$ | Background level of inherent information | 1 | [10] |
| SL | Source level of whale call (dB re 1 m) | 178 | [13] |
| RL<sub>min</sub> | Minimum detectable call (dB re 1 m) | 88 | N/A |
| $N_s$ | Noise avoidance transition parameter | 0.5 | N/A |
| $N_{\text{threshold}}$ | Noise avoidance threshold (dB re 1 m) (Fig. 2,4) | 120 | N/A |
| $N_{\text{threshold}}$ | Noise avoidance threshold (dB re 1 m) (Fig. 3) | 105, 110, 115 | N/A |
| $N_{\text{threshold}}$ | Noise avoidance threshold (dB re 1 m) (Fig. 5, 6) | 115 | N/A |
| $d_{\text{threshold}}$ | Land avoidance threshold (m) | 30 | N/A |
| SNR | Signal-to-noise ratio for detectable calls (dB re 1 m) | -5 | N/A |
| $\gamma$ | Logarithmic sound transmission decay parameter | 17.8 | [12] |
| $d_{\text{target}}$ | Distance to target to be counted as arrived (m) | 50,000 | N/A |
| $\mathbf{x}_{\text{target}}$ | Target location | 61° N, 5° W | N/A |
| $\mathbf{x}_{\text{initial}}$ | Initial location (Fig. 2-4) | 53.5° N, 4.5° E | N/A |
| $\mathbf{x}_{\text{initial}}$ | Initial location (Fig. 5) | 55° N, 4.5° E | N/A |
| $\mathbf{x}_{\text{spread}}$ | Spread in initial location | 1° N, 1° E | N/A |
| SL<sub>construction</sub> | Source level of construction activity (dB re 1 m) | 210 | [9] |

For the results in Figure 5 we generate noise maps corresponding to three different shipping scenarios. The first scenario we consider is shipping traffic that generates migratory behaviour that is similar to that seen for the current soundscape data. We choose to replicate this to ensure that there are not discrepancies between directly compared scenarios due to different choices in sound transmission models. That is, the current soundscape data relies on a partial differential equation model of sound transmission rather than the logarithmic model considered here. The second scenario we consider is a 50% increase in shipping traffic. Specifically, we maintain the same routes, but for each route we increase the number of ships by 50%. The third scenario we consider is a 50% increase in shipping traffic (relative to current) but with sound mitigation measures applied via a slowdown. The ships emit a lower level of noise but require longer to traverse a shipping route. It is of interest to determine whether this may ameliorate the impact of the presence of additional ships. We consider 13 shipping routes (each of which can be traversed forward or backward, for a total of 26 routes). The precise start points, end points and waypoints can be found in the code repository; we also present a representative image that contains this information in Fig. 5 (main manuscript).

### Construction model

To investigate the impact of potential future construction activity on whale migration we examine the introduction of an intermittent noise source. This noise source is considered to be a representation of the noise emitted during pile driving activity in the construction of, say, an oil rig or an offshore wind farm. The source level of the construction activity, while active, is chosen to be 210 dB (re 1 m), consistent with observations [9]. The location of the construction activity is chosen to be at 56.35 *^◦^*N, 4.25 *^◦^*E, though we comment this location is for illustrative purposes only, rather than reflecting any specific construction plans. We consider a range of scenarios involving different levels of activity. These include patterns of work of (i) no activity; (ii) 5 hours of activity, followed by 19 hours of inactivity; (iii) 8 hours of activity, followed by 16 hours of inactivity, and; (iv) 12 hours of activity, followed by 12 hours of inactivity. We examine the situation where this construction activity is occurring in either a pristine soundscape (Fig. 5) or the current soundscape (Fig. 4). We also examine the impact of redistributing the times of activity while maintaining the total activity. Here we consider (i) 6 hours of activity, followed by 18 hours of inactivity, every day; (ii) 9 hours of activity, followed by 15 hours of inactivity, for two days followed by a day of no activity, and; (iii) 12 hours of activity, followed by 12 hours of inactivity, followed by a day of no activity. The results are presented in Figure 5. Note that there is, on average, 6 hours of construction activity each day. In each simulation we randomly select which day of construction activity corresponds to the first day in the simulation. We observe extremely similar results for each construction schedule, which suggests that the total amount of construction activity has a more significant impact than the distribution of the activities.

### Sensitivity analysis

We conduct a sensitivity analysis to examine how the median arrival time of the population varies with changes in the model parameters, with a particular focus on model parameters that are not readily able to be estimated. We first consider changes to the level of inherent information available to individuals and present the median arrival time for the population in the pristine and current soundscapes in Figure 6. We observe a monotonic, but plateauing, decrease in median arrival time with an increase in the inherent information. This is consistent with the results presented in Johnston and Painter [10]. Further, the results indicate that the population is not at risk at migration failure due to the ocean currents overcoming effective navigation ability for the parameter values selected in the main manuscript.

We next consider changes to the initial spacing of the population and present the results in Figure 7. We observe that the median arrival time is insensitive to the initial spread of the population. This is partially due to the fact that the bathymetry can induce aggregation through avoidance of regions of shallow water.

**Figure 1:**
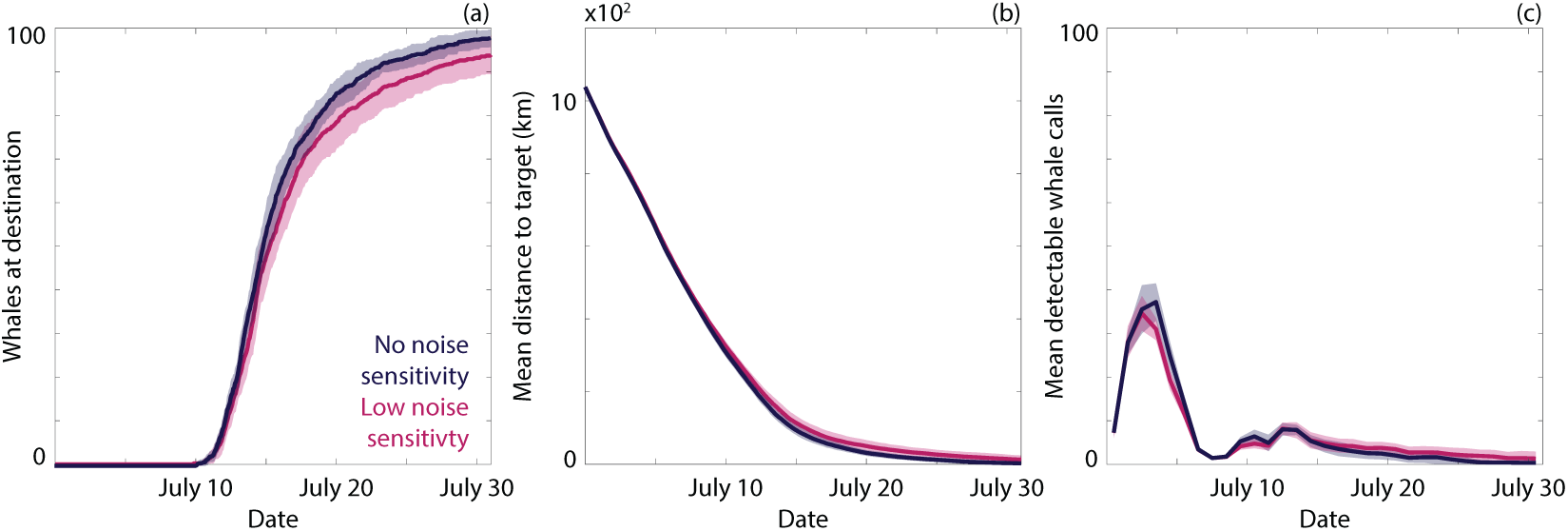
Comparison of navigation with different sensitivities to noise that induce an avoidance response. (a) The number of whales that have arrived at the target destination for no noise sensitivity (dark blue) and low noise sensitivity (magenta). (b) The mean distance of the population from the target for no noise sensitivity (dark blue) and low noise sensitivity (magenta). (c) The mean number of detectable whale calls (averaged daily) for no noise sensitivity (dark blue) and low noise sensitivity (magenta). The lines and ribbons correspond to the mean *±* one standard deviation over 10 simulations.

**Figure 2:**
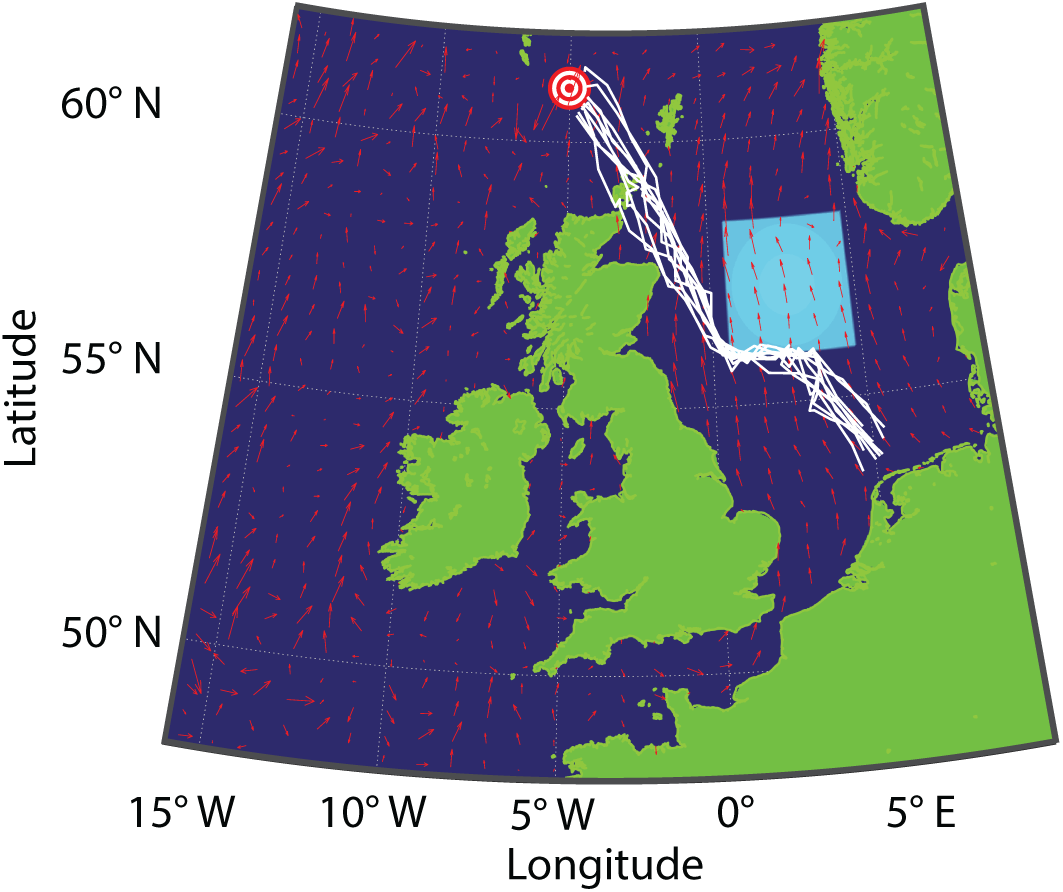
Demonstration of noise avoidance. Representative trajectories of whales avoiding an extreme noise source (cyan) located in the middle of a migration path.

**Figure 3:**
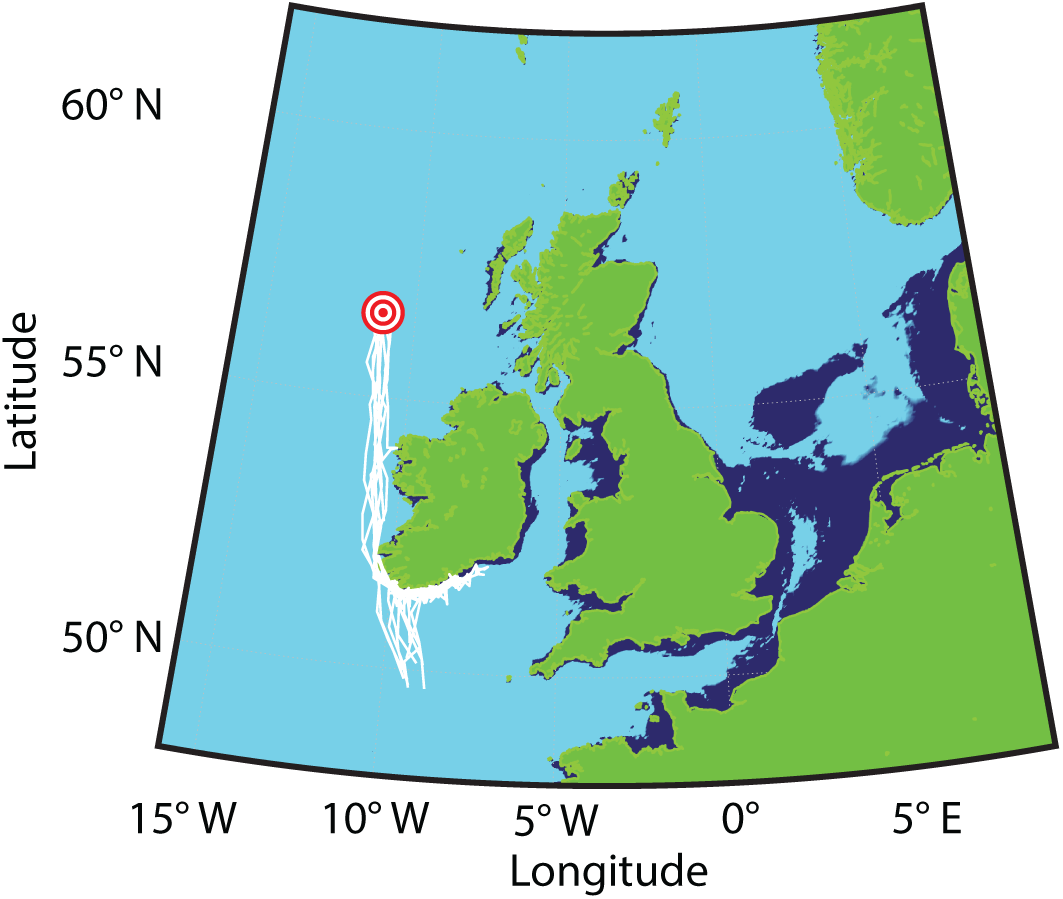
Demonstration of land avoidance. Representative trajectories of whales avoiding land located in the middle of a migration path. Cyan regions correspond to ocean depths of greater than 40m, dark blue regions correspond to ocean depths of less than 40m.

**Figure 4:**
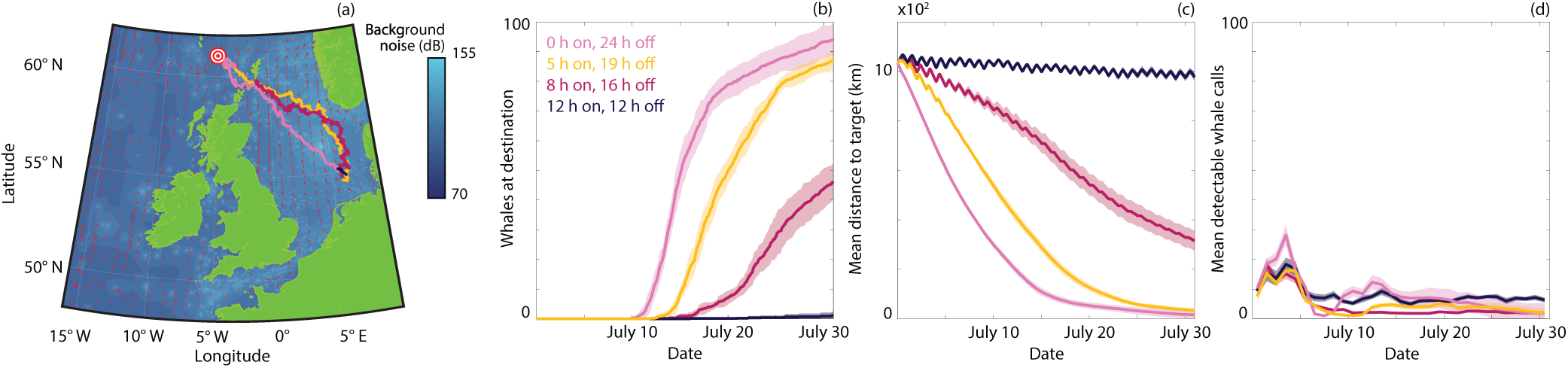
Comparison of navigation under different levels of construction activity in the current soundscape. (a) Noise map during active construction at 56.35 *^◦^*N, 4.25 *^◦^*E with median trajectories for 0 hours (pink), 5 hours (orange), 8 hours (magenta) or 12 hours (dark blue) of construction activity per day. (b) The number of whales that have arrived at the target destination with 0 hours (pink), 5 hours (orange), 8 hours (magenta) or 12 hours (dark blue) of construction activity per day. (c) The mean distance of the population from the target for 0 hours (pink), 5 hours (orange), 8 hours (magenta) or 12 hours (dark blue) of construction activity per day. (d) The mean number of detectable whale calls (averaged daily) for 0 hours (pink), 5 hours (orange), 8 hours (magenta) or 12 hours (dark blue) of construction activity per day. The lines and ribbons correspond to the mean *±* one standard deviation over 10 simulations. In all simulations there are 100 whales.

**Figure 5:**
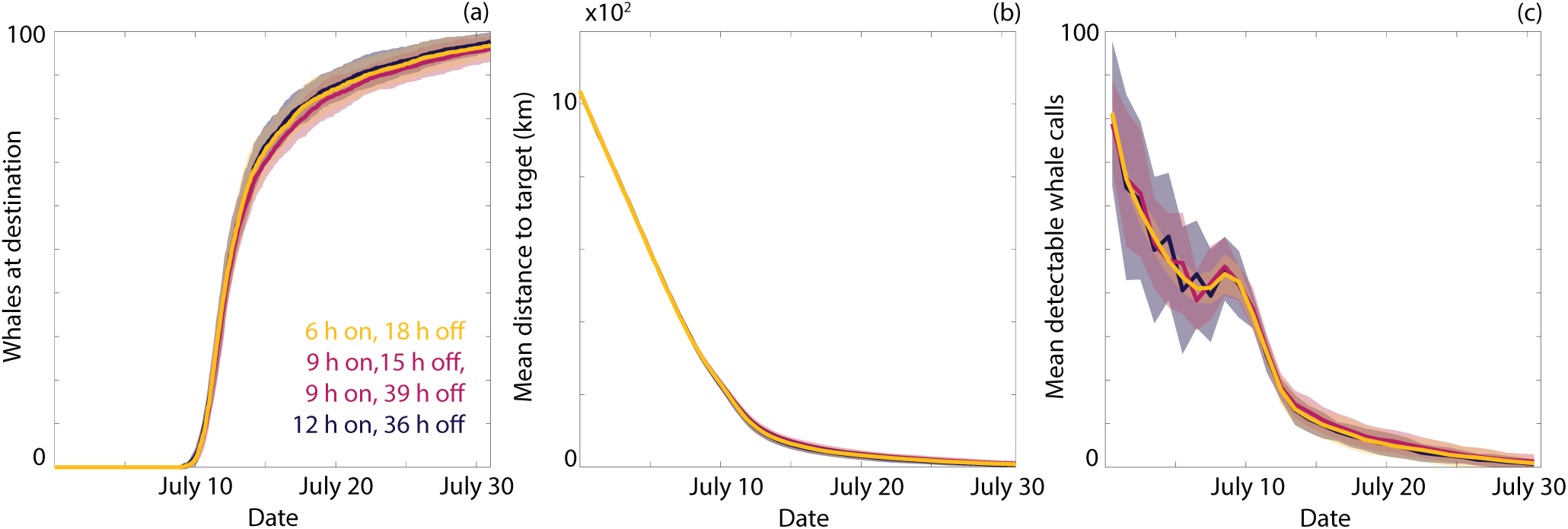
Comparison of navigation under different schedules for construction activity in the pristine soundscape. (a) The number of whales that have arrived at the target destination with 6 hours daily (orange), 9 hours for two days, followed by an off day (magenta), 12 hours for one day, followed by an off day (dark blue) of construction activity. (c) The mean distance of the population from the target with 6 hours daily (orange), 9 hours for two days, followed by an off day (magenta), 12 hours for one day, followed by an off day (dark blue) of construction activity. (d) The mean number of detectable whale calls (averaged daily) with 6 hours daily (orange), 9 hours for two days, followed by an off day (magenta), 12 hours for one day, followed by an off day (dark blue) of construction activity. The lines and ribbons correspond to the mean *±* one standard deviation over 30 simulations. In all simulations there are 100 whales.

**Figure 6:**
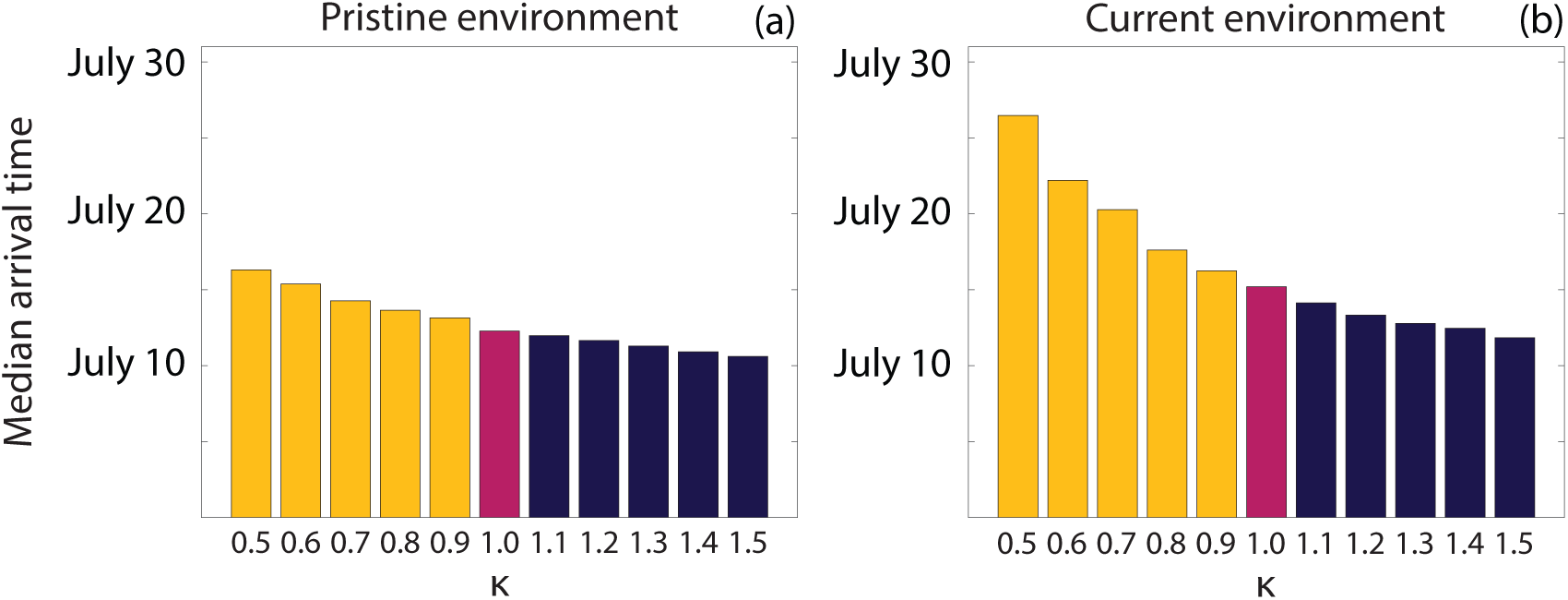
Sensitivity of migration timing to inherent information. Median arrival time for a population of 100 whales for different levels of inherent information in (a) the pristine soundscape and (b) the current soundscape. Results are the mean of 10 identically-prepared realisations of the simulation. Magenta indicates the parameter value used in the main manuscript, orange indicates parameter values that might be expected to delay arrival times, dark blue indicates parameter values that might be expected to expedite arrival times.

**Figure 7:**
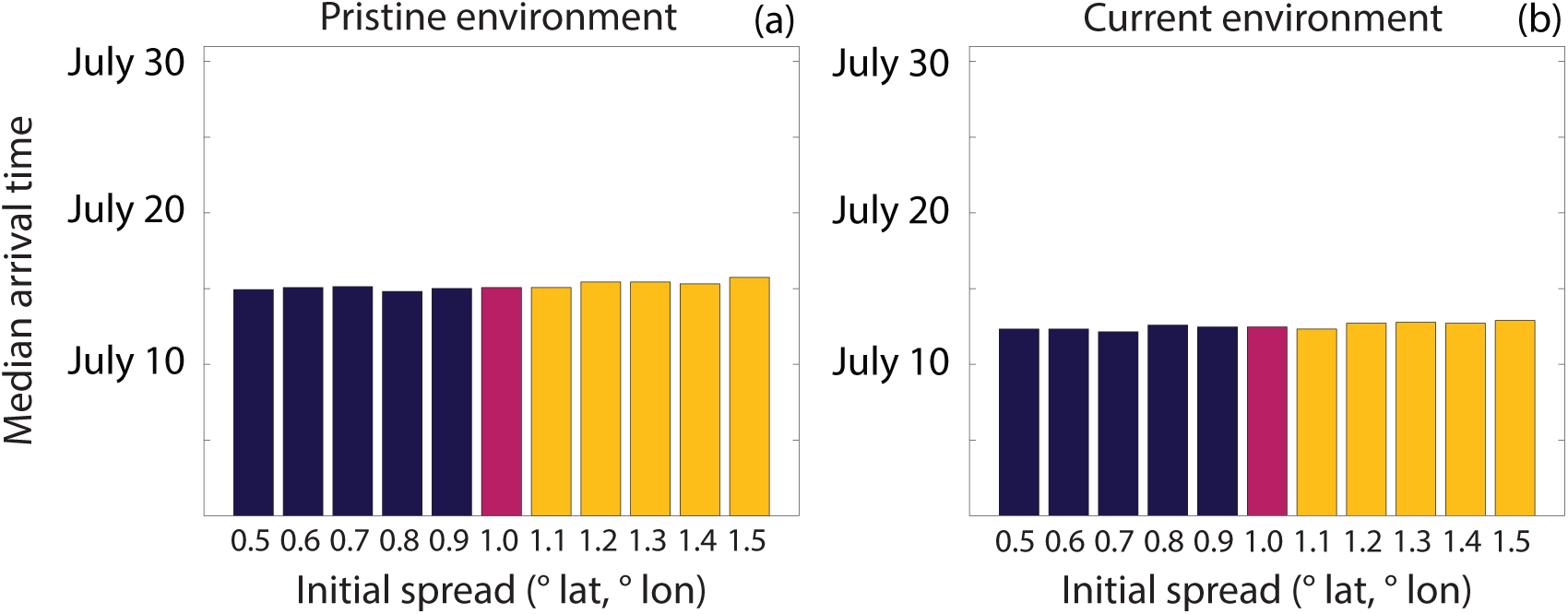
Sensitivity of migration timing to the initial spread of the population. Median arrival time for a population of 100 whales for different levels of inherent information in (a) the pristine soundscape and (b) the current soundscape. Results are the mean of 10 identically-prepared realisations of the simulation. Magenta indicates the parameter value used in the main manuscript, orange indicates parameter values that might be expected to delay arrival times, dark blue indicates parameter values that might be expected to expedite arrival times.

**Figure 8:**
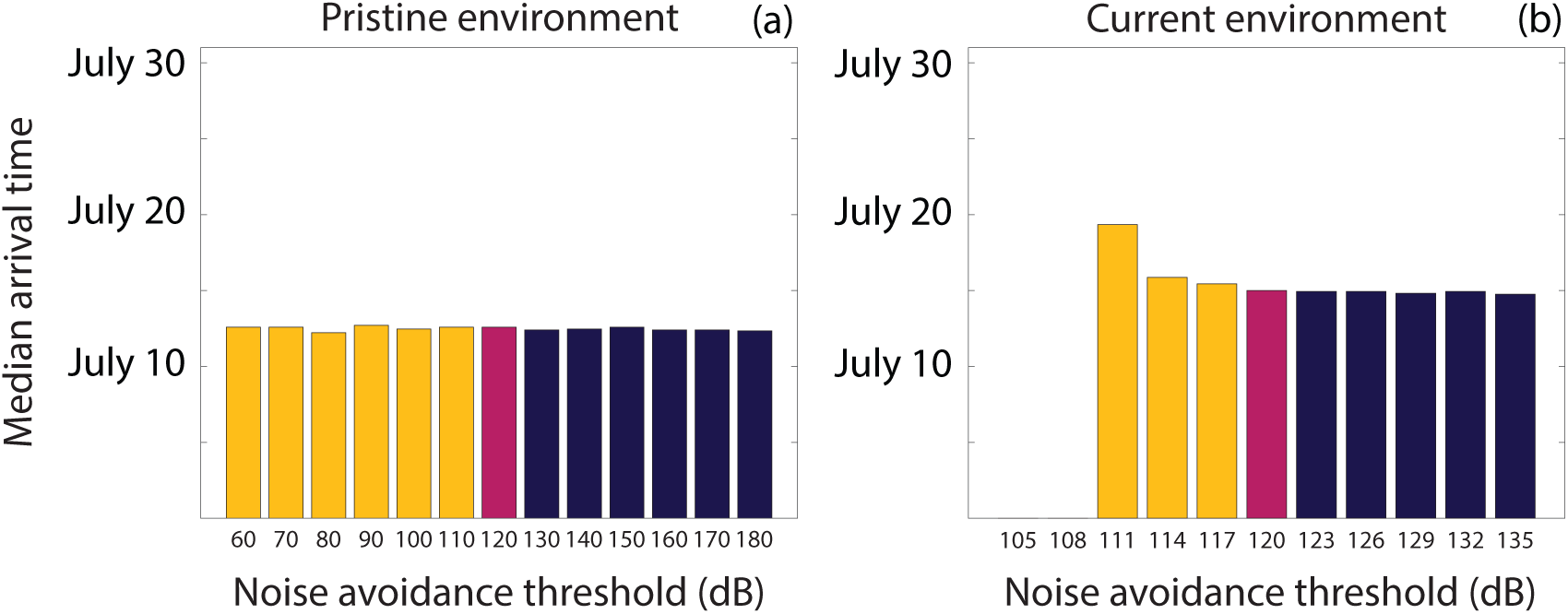
Sensitivity of migration timing to the noise avoidance threshold. Median arrival time for a population of 100 whales for different levels of inherent information with (a) low sensitivity to noise and (b) intermediate sensitivity to noise. Results are the mean of 10 identically-prepared realisations of the simulation. Magenta indicates the parameter value used in the main manuscript, orange indicates parameter values that might be expected to delay arrival times, dark blue indicates parameter values that might be expected to expedite arrival times.

**Figure 9:**
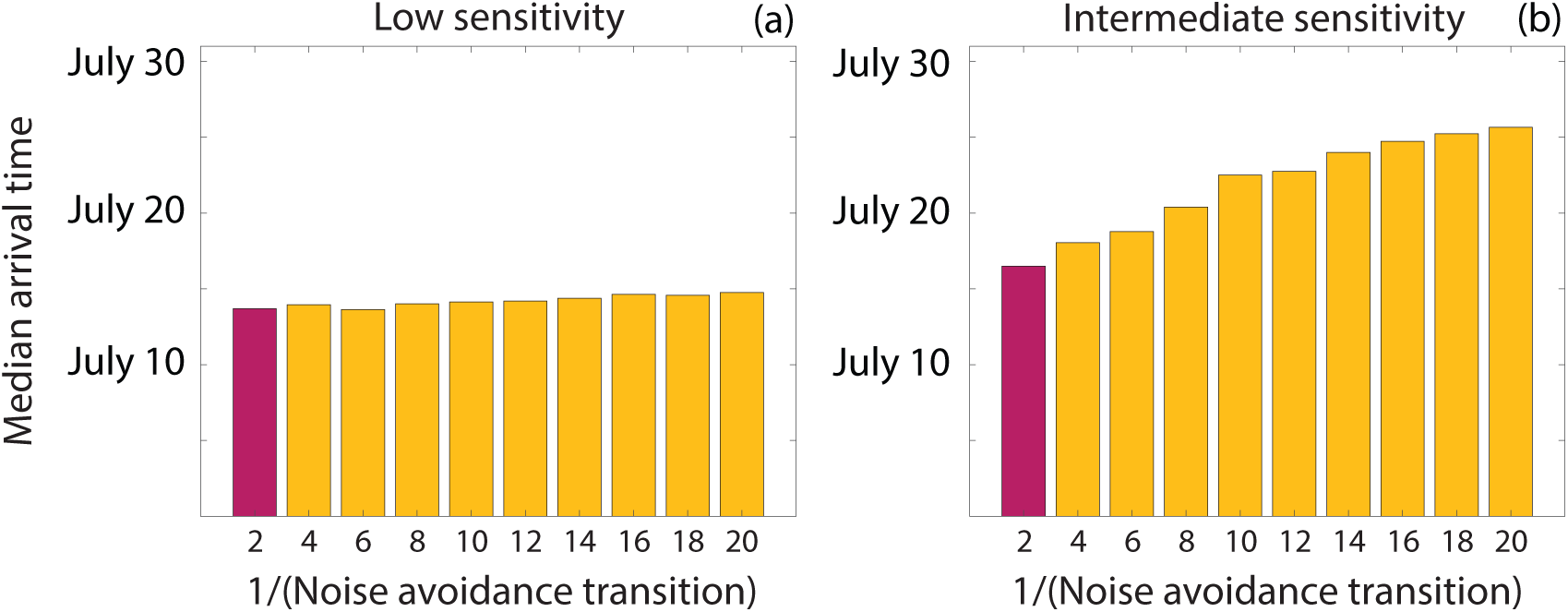
Sensitivity of migration timing to the noise avoidance transition parameter. Median arrival time for a population of 100 whales for different levels of inherent information in (a) the pristine soundscape and (b) the current soundscape. Results are the mean of 10 identically-prepared realisations of the simulation. Magenta indicates the parameter value used in the main manuscript and orange indicates parameter values that might be expected to delay arrival times.

We now consider the influence of the noise avoidance threshold parameter. We perform simulations in the pristine and current soundscapes and present the results in Figure 8. Unsurprisingly, for the pristine sound-scape, the median arrival time is insensitive to a wide range of noise avoidance threshold parameter values as there are very few regions where noise avoidance is relevant for the parameters considered. In contrast, for the current soundscape, the median arrival time is highly sensitive. For noise avoidance threshold parameter values that are well above the background noise, we see no change to the median arrival time. However, there is a rapid transition between a consistent median arrival time and a complete failure of migration. This highlights the need to understand the response of whales to extreme noise sources.

Finally, we consider the how the rate of transition between noise avoidance behaviour and regular migration behaviour impacts the median arrival time. We present the results in Figure 9. A decrease in the rate of transition implies that there is weak, but increased, noise avoidance behaviour at lower noise levels, while there is reduced noise avoidance behaviour at higher noise levels (relative to the original rate of transition). We observe that as the transition becomes slower, the median arrival time increases. This is because it is more common for there to be a non-negligible component of behaviour that is dictated by noise avoidance, compared to a sharp transition. However, experimental observations indicate that this type of slow transition is less likely as there appears to be a threshold noise level below which cetaceans do not avoid the noise source.

## Overview, design concepts and details protocol for

The model description follows the ODD (Overview, Design concepts, Details) protocol for describing individual- and agent-based models [2], as updated in [3].

### Purpose

The purpose of this model is to explore the interaction between noise pollution, and migrating and navigating whale populations. Specifically, we seek to investigate how different potential noise responses at the local scale manifest as different population behaviour (with respect to migration). We consider three mechanisms: noise avoidance, where whales move away from regions of noise above a specified threshold; information loss, where whales lose inherent navigation information due to noise pollution, and; loss of communication space, where the presence of noise pollution reduces the range over which whales can com-municate.

### Pattern

The model should provide results where a loss of either group information (due to reduced communication) or inherent information (due to a reduced ability to detect a navigation cue) at the individual scale should result in a slower rate of migration; consistent with previous models of collective navigation [4]. This is measured through the number of individuals that have arrived at a target destination over time and the average distance between the population and the target destination.

### Entities

The model contains the following entities: individual agents that represent whales, and a global environment that contains information about, for example, the local noise level, ocean currents, and bathymetry. Table 1 contains the list of variables in the global environment, alongside their units and meaning. Table 2 contains the list of state variables for the agents, alongside their units and meaning.

### Scales

In all cases, the model is performed on a two-dimensional continuous space simulation domain from 48 *^◦^*N to 62 *^◦^*N in latitude and from 16 *^◦^*W to 9 *^◦^*E in longitude. Note that this is an area coinciding with known distributions of baleen whale populations. To be consistent with the external environment data, we impose a discretisation such that there are 281 discrete latitude points and 301 discrete longitude points for any global environment data. Agents exist in continuous space. The model runs over the course of a single month (though we note that this is trivial to change by changing *tEnd*). This month is nominally July due to the selected global environment data. Time evolves in a continuous manner. No other dimensions are represented.

**Table 1:** List of variables that are components of the global environment.

| Variable name | Variable type and units | Meaning |
| --- | --- | --- |
| *backgroundNoise* | Real number (dB re 1m) | Ambient noise at location in time and space. |
| *depthGrid* | Real number (m) | Depth of the ocean at each location |
| *latFlowVelocity* | Real number (m/h) | Ocean current velocity in lat. direction at location in time and space. |
| *lonFlowVelocity* | Real number (m/h) | Ocean current velocity in lon. direction at location in time and space. |
| *meanWindNoise* | Real number (dB re 1m) | Mean wind noise at location in space. |
| *shippingNoise* | Real number (dB re 1m) | Ambient shipping noise at location in time and space. |
| *navigationField* | Real number | Direction of target destination at location in space. |
| *navigationStrengthField* | Real number | Level of inherent information about target destination at location in time and space. |

**Table 2:** List of state variables for the agents.

| Variable name | Variable type and units | Meaning |
| --- | --- | --- |
| *alpha* | Real number, static | Weighting of inherent and group information for heading. |
| *avoidNoiseWeight* | Real number, variable | Strength of noise avoidance response. |
| *avoidShallowWaterWeight* | Real number, variable, | Strength of shallow water avoidance response. |
| *backgroundNoiseAtLocation* | Real number, variable (dB re 1m) | Level of ambient noise at current location. |
| *bestGuessHeading* | Real number, variable | Estimate of target destination direction. |
| *bestGuessStrength* | Real number, variable | Estimate of confidence in target destination direction. |
| *beta* | Real number, static | Weighting of inherent and group information for confidence. |
| *communicatedDirection* | Real number, variable | Communicated direction of target destination. |
| *depthAtLocation* | Real number, variable (m) | Depth of water at current location. |
| *directionAwayFromNoise* | Real number, variable | Direction of maximal decrease in ambient noise at current location. |
| *directionOfDeepestWater* | Real number, variable | Direction of maximal increase in water depth at current location. |
| *distanceFromGoal* | Real number, variable (m) | Distance between the agent and the target destination. |
| *heading* | Real number, variable | Current heading of the agent. |
| *initialPosition* | Real number, static | Initial location of the agent. |
| *latPosition* | Real number, variable | Current latitude position of the agent. |
| *lonPosition* | Real number, variable | Current longitude position of the agent. |
| *minimumHearing* | Real number, static (dB re 1m) | Minimum noise signal that can be detected by an agent. |
| *minimumNoiseOverlap* | Real number, static (dB re 1m) | Signal-to-noise ratio for signal detection. |
| *neighbours* | Integer vector, variable | List of other agents detectable by the agent. |
| *nNeighbours* | Integer number, variable | Number of other agents detectable by the agent. |
| *noiseAvoidanceLevel* | Real number, static (dB re 1m) | Threshold noise level for the noise avoidance response. |
| *position* | Real vector, variable | Current $(x, y)$ position of the agent. |
| *potentialHeading* | Real number, variable | Direction of target destination from inherent information only. |
| *runTime* | Real number, static (h) | Average time between reorientation events. |
| *timeToUpdate* | Real number, variable (h) | Time until the agent undergoes a reorientation event. |
| *velocity* | Real number, static (m/h) | Swimming speed of an agent. |

### Processes

The model describes the migration of a whale population that is initially located in the south of the North Sea with a target destination north of Scotland. The main components of the model are the “run” and “tumble” phases of a velocity jump random walk. The run component simply consists of ballistic motion of agents in the direction selected in the previous tumble phase combined with motion driven by immersion in ocean currents. The tumble phase is a reorientation event where inherent information, group information, land avoidance behaviour and noise avoidance behaviour are all combined to select a new heading. Each component of the reorientation event is described in the sub-models section below. In the run phase *latPosition*, *lonPosition*, *distanceFromGoal*, and *timeToUpdate* are all changed. In a reorientation event *avoidNoiseWeight*, *avoidShallowWaterWeight*, *backgroundNoiseAtLocation*, *bestGuessHeading*, *bestGuessStrength*, *communicatedDirection*, *depthAtLocation*, *directionAwayFromNoise*, *directionOfDeepestWater*, *heading*, *neighbours*, *nNeighbours* and *potentialHeading* are all updated.

### Schedule

The run phase for an agent lasts for an exponentially distributed length of time, with the mean length defined by *runTime*. The tumble phase is instantaneous; a reorientation event occurs at the end of each run phase. As time is continuous and run duration is a random variable, agent reorientation is asynchronous across the population. After one reorientation event, time is simply advanced to the next reorientation event in the population.

### Basic principles

The model seeks to describe the collective navigation of a whale population, where it is feasible that individual whales rely on a sophisticated synthesis of navigation cues and information from con-specifics. The broader problem can be considered as an example of the “wisdom of the crowd” phenomenon (see [1], for example), where groups are more effective at migrating than individuals. This specific model extends the previous model of Johnston and Painter [4], where confidence or uncertainty in navigation is explicitly included in the information synthesis process. Here the model is tailored to whale populations and sound transmission in the oceans, where communication over long distances is possible under pristine ocean conditions, and the communication space is reduced under the current ocean conditions. We introduce sub-models to describe sound transmission, noise avoidance responses, land avoidance responses and inherent information loss.

### Emergence

The model provides both qualitative and quantitative outcomes. It allows us to quantify the changes in summary statistics, such as the population migration rate and the percentage of successful migrations, due to changes in the global environment or parameters in the model. Moreover, clear qualitative differences in migration patterns arise in response to different types of reactions to high background noise levels. While these patterns rely on the form of the reaction (avoidance, loss of communication, loss of inherent information), these reactions are based on clear physical principles (observed retreating from extreme noise sources, and sound transmission in water).

### Adaption

Agents update a considerable number of state variables (see Table 2) throughout the model simulation. The ultimate decision that is made is the new heading of the agent. This depends on the current location (via the global environmental variables of the ambient noise, bathymetry, navigation field and inherent information) and the neighbours of the agent. This process is described in depth in the reorientation model. An update of *heading* necessarily involves updating *bestGuessHeading*, *bestGuessStrength*, *heading*, *potentialHeading*, *timeToUpdate* and *communicatedDirection* (other state variables are also updated; see discussion of reorientation). The state variables of *distanceToGoal* and *position* changes constantly during the run phase. The reorientation process is not deterministic: the final choice of heading is a sample from a determined probability distribution. This can be considered as an example of indirect objective seeking, as the agents do not always progress towards the target destination (i.e. the objective).

### Objectives

Agents in the model have only a single object: migrate towards a target destination until they are sufficiently close to that destination, upon which migration is classified as successful. This is tracked through the *distanceToGoal* state variable, which simply calculates the Euclidean distance between the agent and the target destination.

### Learning

This model does not include learning behaviour.

### Prediction

The model includes implicit prediction in both noise and land avoidance behaviour. We choose to model this behaviour via taxis-type responses, where we assume the individual can estimate or detect the spatial change in noise and ocean depth, and move down or up this gradient as appropriate. Many animals are known to move via taxis-type responses to a cue and hence it is a suitable modelling choice that will approximate known behaviour in the absence of detailed observations. There is an implicit assumption that the gradient can be estimated; however, this only relies on the ability to detect a cue nonlocally (but not over a significant distance) or remember a level of a cue at a previous location, which is plausible for whales.

### Sensing

Agents are able to use all of their state variables in their decision making. The updating of the state variables involved in navigation includes uncertainty. As such, while the agents are able to reliably use state variables, the process of obtaining the state variables is where the uncertainty manifests itself. For example, *bestGuessHeading*, *bestGuessStrength*, *heading*, *potentialHeading*, *timeToUpdate* and communicatedDirection are all updated according to uncertain information. The distance over which an agent can detect other agents is a function of the local environment and the source level of the communication of the other agents.

### Interaction

Interaction between agents is indirect and solely based on communication. Agents emit calls at a specified source level, assumed to be constant across the population. The calls propagate through the ocean environment according to a logarithmic decay model. Agents can detect emitted calls if the received level of the call is within a specified signal-to-noise ratio of the local background noise. The mathematical description of the sound propagation is defined in the sub-models section below. Interactions between agents and the environment is a purely local interaction.

### Stochasticity

There are four forms of stochasticity in the model. First, the inital locations of individuals (*initalPosition*) are uniformly randomly distributed within a specified area of space. Second, the duration of each run phase (*timeToUpdate*) for each individual is sampled from an exponential distribution with mean parameter *runTime*. Third, the individual’s estimate of the target destination (*potentialHeading*) is sampled from a von Mises distribution centred at the target destination direction with concentration parameter that is equal to the level of inherent information. Finally, the individual’s heading (*heading*) is sampled from a von Mises distribution centred at *bestGuessHeading* with concentration parameter *bestGuessStrength*.

### Collectives

There are emergent collectives of agents in the model. The reorientation process is a function of the detected headings from other agents. As such, if there are agents in close proximity, the decision making process indirectly changes. However, this is not explicitly modelled via a collective entity.

### Observation

We record three metrics from the simulation (noting that others could be easily recorded). First, we record the number of agents that have reached the target destination by each time point. This is recorded at 501 time points over 744 hours (one month). Second, we record the average (Euclidean) distance between the agents and the target destination. This is recorded at 501 time points over 744 hours (one month). Finally, we record the average number of detected agents. This is calculated as a daily average and hence is recorded at 31 time points over 744 hours; the reason for the coarseness in the measurement is due to the noisiness of the data.

### Initialisation

The model is initialised by loading the relevant input data (see Input data section). This includes loading the appropriate suite of parameter values for a specific case study (see Input data section). Observation metrics are initialised for later use. Agents are created, the number of which is defined in the suite of parameter values. The initial locations of agents are sampled from the appropriate uniform distributions. The initial length of the run phases are sampled from the appropriate exponential distribution.

### Input data

The model relies on reading in a number of input datasets. The shipping noise (*backgroundNoise*), the wind noise (*meanWindNoise*), the coastal boundary data (*coarseLandX*, *coarseLandY*, *coarseLat*, *coarseLon*), the ocean current data (*latFlowVelocity*, *lonFlowVelocity*) and the look-up table for the von Mises likelihood (*kappaCDF*) are all loaded at the start of the simulation. The *backgroundNoise* may come from validated shipping data, synthetic shipping data (see Supplementary Information) or resource extraction platform data (see Supplementary Information). A particular parameter set (the string *parameterSet*) is loaded to describe the relevant case study or example, which contains the parameters defined in Table 3. Certain parameters are universal in all case studies considered here and are hence defined at the beginning of each simulation.

### Sub-models

The key processes in the model are the reorientation events, which are composed of five different sub-models.

The first sub-model is the inherent information sub-model. When *reduceInformation* has a value of ‘no’ then the level of inherent information about the target destination (and hence the concentration parameter in the von Mises distribution) is constant. When *reduceInformation* has a value of ‘yes’ then the level of inherent information about the target destination varies with space and time. To implement this we choose *κ*(x*, t*) as a function of the noise:

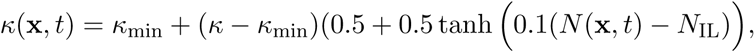

where *κ*_min_ is the minimum level of inherent information (i.e. at extreme noise levels) and *N*_IL_ is a threshold parameter that represents the noise level at which half of the inherent information (that can be lost) is lost. Therefore, the value of *potentialHeading* for an agent relies on the interaction between an agent’s location and this function.

The second sub-model is the sound transmission model. As discussed in the main manuscript, we use a simplified sound transmission model. The received level RL (dB) at a distance *r* (m) from a sound with source level SL (dB), can be modelled by

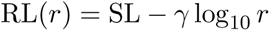

where *γ* log_10_ *r* describes the transmission loss. The coefficient *γ* is bounded below by 10, corresponding to shallow water in which the spreading is effectively cylindrical, and above by 20, for deep water in which sound propagates in all directions (spherical). This sound transmission model allows to calculate which agents can be detected by a specific agent. If RL(*r*) *− N* (x*, t*) *>* SNR, where SNR is the signal-to-noise ratio, for a pair of agents separated by Euclidean distance *r*, then the agents are able to detect each other’s calls. The agents are therefore classed as neighbours. The heading of all neighbour agents contribute to reorientation events and these headings are classified as group information. Full details for the synthesis of group and inherent information is given in [4]. Briefly, the weighted circular mean and resultant vector length are calculated from the heading of an agent and the headings of its neighbours. The weightings are *alpha* and *beta*, respectively. The weighted circular mean gives *bestGuessHeading* while *bestGuessStrength* is obtained from the look-up table *kappaCDF* according to the process described in [4]. The value of *heading* is then sampled from a von Mises distribution with location and concentration parameters *bestGuessHeading* and *bestGuessStrength*.

The third sub-model is the noise avoidance model. To describe noise avoidance behaviour we implement a negative phonotaxis response (i.e. motion in the direction of decreasing noise). The direction of decreasing noise is calculated using a finite difference approximation to the directional derivative from the information stored in *backgroundNoise*. The noise avoidance response is implemented in a weighted manner, where at low noise levels there is essentially no response, and at high noise levels, essentially all motion is determined by the noise avoidance response. Specifically, we calculate the proportion of motion that is driven by noise avoidance *w_na_*(*N* (x*, t*)), where the *N* (x*, t*) is the noise level at location x and time *t*. The remaining proportion of motion (i.e. 1 *− w_na_*(*N* (x*, t*))) is motion corresponding to regular migration behaviour (i.e. in the direction of *heading*). We calculate the weighting via

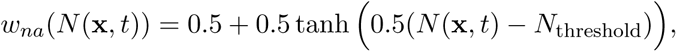

where *N*_threshold_ is a threshold parameter that represents the noise level at which there is an equal weighting between noise avoidance and migration. This weighting is stored as *avoidNoiseWeight*.

The fourth sub-model is the land avoidance model. We consider a similar approach for land avoidance as for noise avoidance, assuming that excessively shallow water will trigger a response in which motion is in the direction of greatest water depth (bathotaxis). As before, this relies on a finite difference approximation of the directional derivative, from the information stored in *depthGrid*. Specifically, we define a weighting *w_la_*(*d*(x)) that represents the proportion of motion that is in the direction of greatest water depth, given the depth at the current location *d*(x). Similar to the noise avoidance response, the remaining proportion of motion (i.e. 1 *− w_la_*(*d*(x))) is motion corresponding to regular migration behaviour (i.e. in the direction of *heading*). We calculate the weighting via

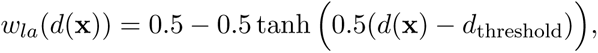

where *d*_threshold_ is a threshold depth that represents the water depth at which there is an equal weighting between land avoidance and migration. This weighting is stored as *avoidShallowWaterWeight*.

Reorientation events therefore occur in the following way. Based on the inherent information model, an initial heading *potentialHeading* is determined. Next, based on the sound transmission model, an agent’s neighbours are calculated. The headings of the neighbours are combined with *potentialHeading* to obtain a preliminary value of *heading*. To account for land avoidance and noise avoidance, *heading* is updated to be a weighted combination of the original *heading* value, and *directionOfDeepestWater* and *directionAwayFromNoise*, with weightings *avoidShallowWaterWeight* and *avoidNoiseWeight*.

The fifth and final sub-model is a failsafe land avoidance model. Here we store information about the location of the coastline in *coarseLandX* and *coarseLandY*. If an agent is determined to have crossed a coastline onto land, the movement is aborted. This is calculated via Matlab’s *inpolygon* function, where the polygons are defined by the coastline points. We use a nested approach for efficiency, first using extremely coarse coastlines stored in *cLX* and *cLY*. If an agent is in these extremely coarse polygons, then we test via less coarse polygons. Given the tortuousity of coastlines, we cannot use fully-detailed coastline data with any reasonable computational efficiency.

**Table 3:**
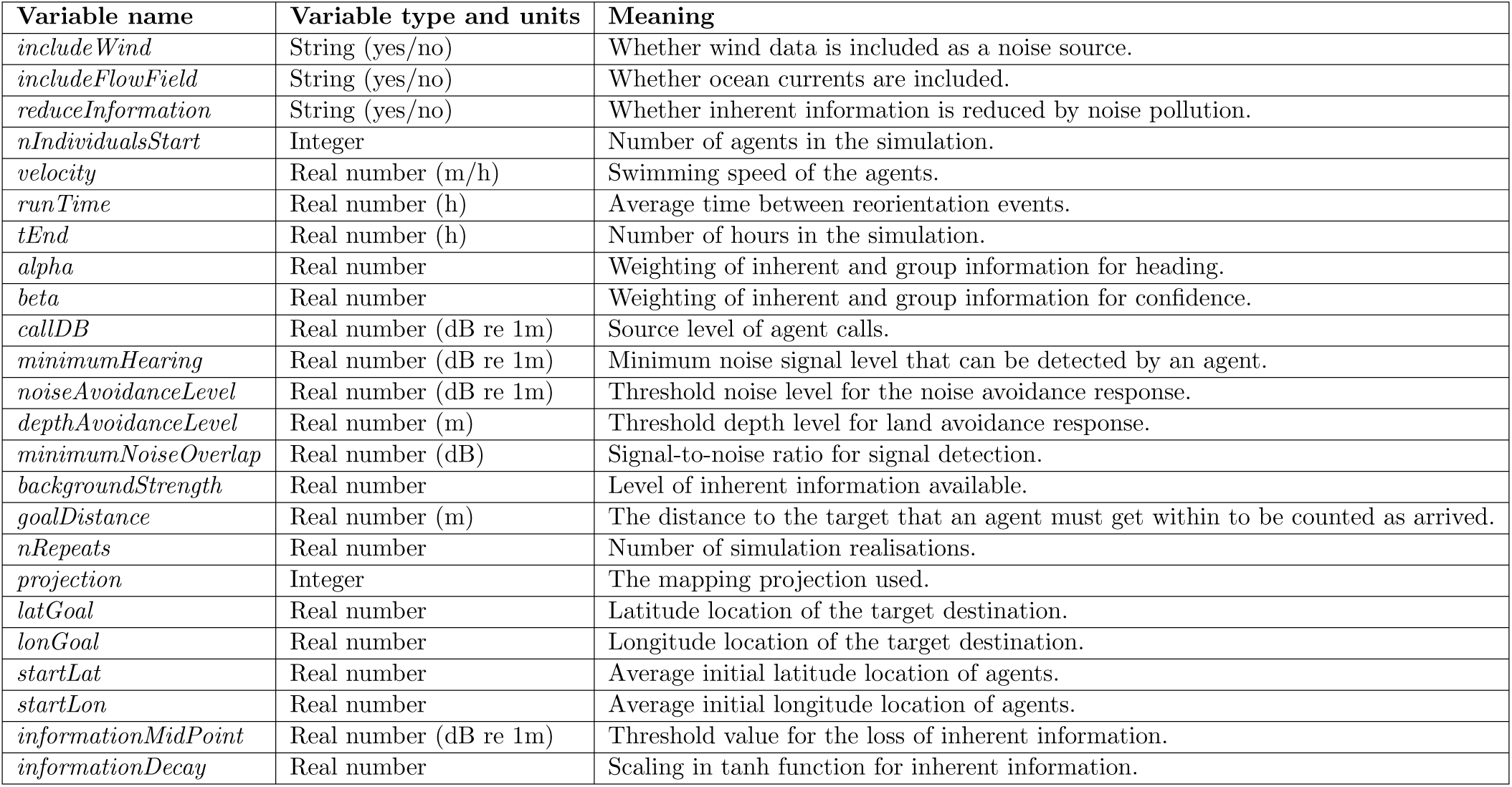
Parameter values loaded during the initialisation process.

